# Comparative Temporal Transcriptomic Analysis of *SOD1* Mutations in iPSC-Motor Neurons

**DOI:** 10.1101/2025.08.19.670632

**Authors:** Miaodan Huang, Ke Zhang, Feng Yu, Qing Chen, Yixia Ling, Pilong Li, Dajiang Qin, Huanxing Su

## Abstract

Mutations in the *SOD1* gene are among the most significant genetic contributors to amyotrophic lateral sclerosis (ALS), with different variants linked to varying disease severity. To investigate the molecular mechanisms driving this variability, we conducted RNA sequencing on spinal motor neurons (MNs) differentiated from isogenic human induced pluripotent stem cell (iPSC) lines engineered via CRISPR/Cas9. These lines carried two representative *SOD1* heterogenous mutations, D91A and G94A, and were analyzed at Days 10 and 20 of neuronal maturation stage to capture the temporal changes of gene expression. We aim to explore how these mutations affect MN function, identify distinct molecular pathways that may explain the variable severity of ALS, and investigate the role of translation and metabolic dysregulation in disease progression.

## Introduction

Amyotrophic lateral sclerosis (ALS) is a neurodegenerative disease characterized by the progressive degeneration of motor neurons (MNs), leading to muscle weakness, paralysis, and a median survival of 2-4 years (Feldman et al. 2022). The disease remains incurable. Current therapies, such as riluzole (Dharmadasa and Kiernan 2018) and edaravone (Rothstein 2017), primarily target oxidative stress pathways and offer only limited survival benefits.

Mutations in the *SOD1* gene are among the most common genetic causes of ALS. Recently, antisense oligonucleotide (ASO) therapies targeting *SOD1* mRNA have emerged as promising treatments by lowering SOD1 protein levels and slowing disease progression in *SOD1*-associated cases (Miller et al. 2020; Miller et al. 2022; Blair 2023). However, most *SOD1*-associated ALS patients are heterozygous, and current ASOs reduce both mutant and wild-type (WT) alleles, which may unintentionally compromise the physiological role of WT SOD1, exacerbating oxidative stress and cellular dysfunction while overlooking potential mutation-specific pathogenic mechanisms.

Our previous study (Huang et al. 2024) demonstrated that distinct *SOD1* mutations are associated with variability in geographic patterns, clinical heterogeneity, disease severity, and molecular alterations, underscoring the importance of mutation-specific mechanisms in ALS pathogenesis. Given this heterogeneity, therapeutic strategies such as ASOs should not only target mutant *SOD1* expression but also consider mutation-specific cellular dysfunctions that drive disease progression. Moreover, as ALS typically manifests with late onset but rapid progression, elucidating the molecular basis of this temporal pattern is critical for understanding disease dynamics and improving therapeutic design.

In this study, we investigated the molecular consequences of *SOD1* mutations by transcriptomic profiling of induced pluripotent stem cell (iPSC)-derived MNs. By comparing the effects of two representative *SOD1* variants, D91A and G94A, at two different time points, we aim to uncover mutation-specific temporal pathological mechanisms at early disease stages and provide mechanistic insights that may inform the development of more precise and integrative therapeutic strategies.

## Results

### Establishment of Isogenic iPSC Lines carrying different *SOD1* Genotypes

To dissect mutation-specific mechanisms in *SOD1*-associated ALS, we selected two representative mutations: D91A, the most prevalent *SOD1* variant in ALS considering both heterozygous and homozygous cases, and G94A, a well-studied mutation associated with established ALS mouse models (Gurney et al. 1994). Using CRISPR/Cas9 RNP-mediated genome editing (Okamoto et al. 2019), we generated isogenic iPSC lines carrying WT/WT and WT/G94A genotypes from a parental iPSC line originally derived from a female ALS patient heterozygous for the D91A mutation (WT/D91A) (Supplemental Fig 1A). A guide RNA (gRNA) specifically targeting the D91A allele was co-delivered with single-stranded oligodeoxynucleotide (ssODN) templates (Supplemental Table 1) to either correct the mutation to WT or introduce the G94A mutation.

High editing efficiencies were achieved, with ICE analysis inferring 96% indels for D91A correction and 36% for G94A introduction (Supplemental Fig 1B). After accounting for the monoallelic targeting of the D91A haplotype, the calculated efficiencies were 92% and 72%, respectively. Precise editing was confirmed in 13 of 15 colonies for D91A correction and 36 of 49 clones for G94A introduction, yielding on-target success rates of 86.67% and 73.47%, respectively, closely matching the calculated efficiencies. The high editing efficiency likely resulted from the specific gRNA targeting of the D91A allele, sparing the WT copy and preventing repeated cleavage. In the G94A-edited lines, the mutation altered the PAM sequence from TGG to TGC, which may have further enhanced editing efficiency.

To assess the genomic integrity, we performed karyotype analysis, off-target PCR, and teratoma assays on representative clones from each editing strategy (Supplemental Fig 1C-D). Karyotype analysis showed no chromosomal abnormalities, indicating genomic stability. The teratoma assays demonstrated the capacity for differentiation into all three germ layers, confirming preserved pluripotency. Sanger sequencing at 16 predicted off-target sites (Supplemental Table 2) detected no off-target effects, revealed no unintended genomic alterations.

To characterize the SOD1 protein variants expressed from the edited alleles, we performed western blot analysis to examine their electrophoretic mobility. The WT/D91A variant exhibited two distinct bands: one corresponding to the migration patterns of WT/WT and WT/G94A, and a second band that migrated more rapidly (Supplemental Fig 1E). This observation aligned with previous reports showing that the D91A mutant exhibited faster migration than WT SOD1 due to the a reduced molecular weight caused by the Asp-to-Ala substitution (Marklund et al. 1997), whereas the G94A mutant migrated similar to WT (Hayward et al. 2002), confirming successful genome editing and the presence of the expected protein variants.

Furthermore, immunofluorescence analysis showed consistent expression of key pluripotency markers, including OCT4, SOX2, SSEA4, and NANOG, across all edited and parental lines (Supplemental Fig 1F-G), confirming the maintenance of pluripotent identity.

In conclusion, we established a robust platform of isogenic iPSC lines carrying WT/WT, WT/D91A, and WT/G94A *SOD1* genotypes with identical genetic background while preserving genome integrity and normal pluripotency.

### Evaluation of the identity of MNs Differentiation from iPSC Lines

To elucidate the biological effects of *SOD1* mutations associated with ALS, we differentiated our established iPSC lines into spinal MNs (Fig. 1A) following the published protocol by Du *et al*.(Du et al. 2015) with a few modifications. To assess disease phenotypes, MNs were cultured without neurotrophic factors. The differentiated cells exhibited characteristic neuronal morphology, with comparable differentiation efficiency across the three cell lines (Fig. 1B).

**Figure 1.**
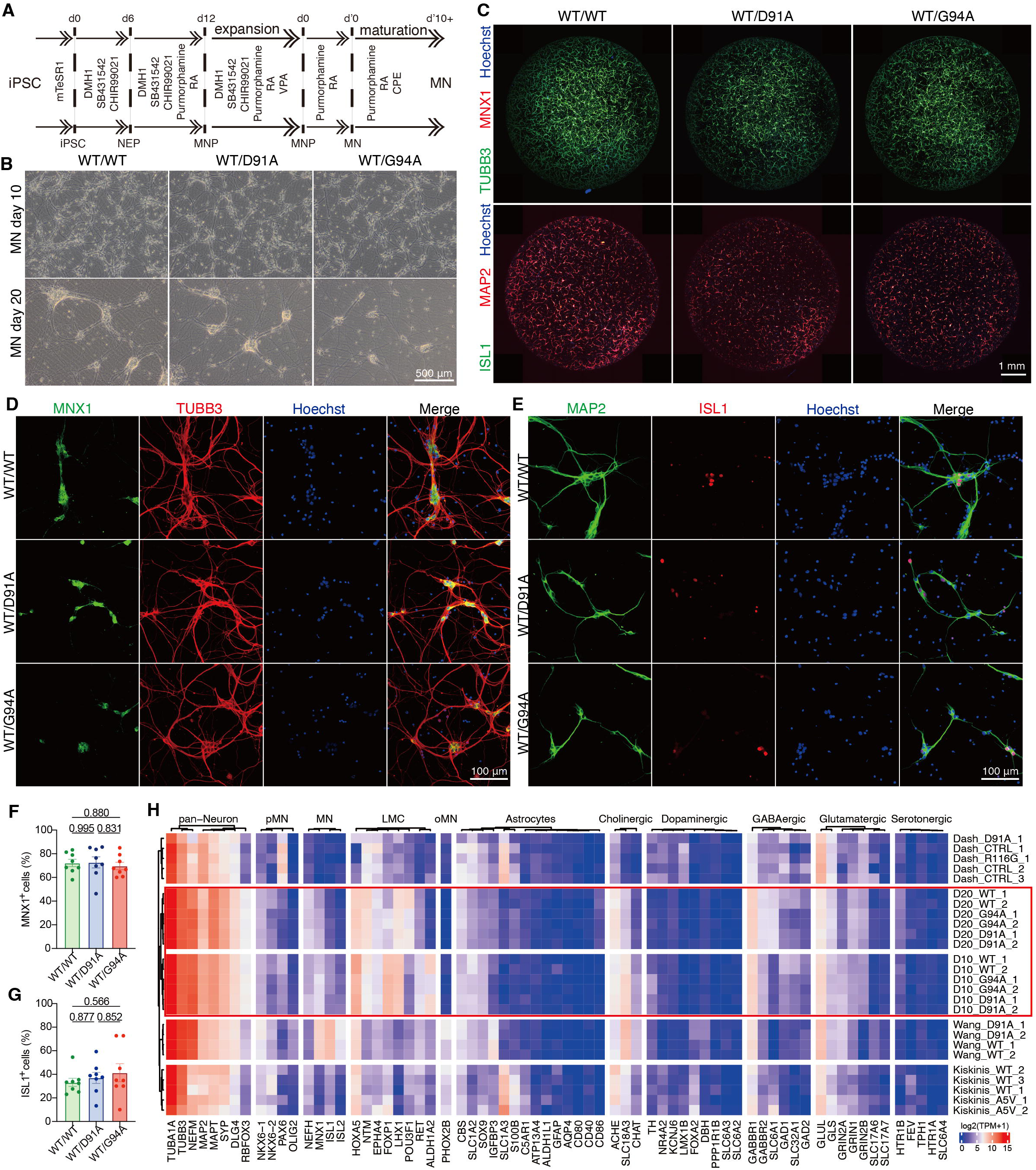
Evaluation of the identities of MNs differentiated from iPSC. (**A**) Diagram of the MN differentiation protocol. (**B**) Bright field microscopy images of MNs at Day 10 and Day 20 of differentiation. (**C-E**) Immunofluorescence analysis of MN markers on Day 10 using primary antibodies against MNX1, ISL1, TUBB3, and MAP2. Images were captured with an Olympus IXplore SpinSR microscope. **C** shows a whole well of a 96-well plate; **D** and **E** were captured in HDR mode. (**F-G**) Proportions of MNX1_ and ISL1_ cells normalized to live Hoechst_nuclei. Data from four fields per well across two wells (n = 8 per condition). (**H**) RNA-seq based expression (Log_(TPM + 1)) of markers for various neuronal subtypes across datasets.

Despite efforts to recapitulate embryonic neurodevelopment, iPSC-derived MN cultures often remain heterogeneous (Giacomelli et al. 2022). To quantify the proportion of MNs in our cultures, we performed immunofluorescence staining for pan-neuronal markers TUBB3 and MAP2, as well as MN-specific transcription factors MNX1 (HB9) and ISL1. By Day 10 of MN maturation, nearly all cells expressed TUBB3 and MAP2 (Fig. 1C-E). The proportions of MNX1-positive cells were comparable across genotypes (72%-73%), while ISL1-positive cells ranged from 33-41% (Fig. 1F-G), with no significant differences among lines. The lower ISL1 percentages suggested some heterogeneity in MN maturation or subtype composition which might result from the lack of neurotrophic factors in maturation stage.

To further assess MN identity and differentiation efficiency for downstream analysis, we examined the expression levels of various cellular markers using bulk RNA-seq and compared them to previously published datasets (Kiskinis et al. 2014; Wang et al. 2017; Baxi et al. 2022; Dash et al. 2022) (Supplemental Fig S1H). Notably, the D91A cell line from Wang *et al*. shares the same genetic background as the WT/D91A iPSC line used in this study, while the corrected lines were independently generated.

We evaluated expression levels of markers including *NEFM*, *MAP2*, *TUBB3*, *TUBA1A*, *MAPT*, *SYP*, *DLG4*, and *RBFOX3* for pan-neuronal characterization. *NEFM* encodes a neurofilament critical for axonal structure; *MAP2* is a dendritic marker; *TUBB3* and *TUBA1A* are essential for neuronal microtubule formation, with *TUBB3* indicating early differentiation and *TUBA1A* reflecting later maturation. MAPT stabilizes microtubules in axons, supporting neuronal integrity and polarity. *SYP* and *DLG4* encode key synaptic proteins, while *RBFOX3*, also known as NeuN, is a well-established marker for mature neurons. We observed high expression levels of these markers in MNs at Days 10 and 20, indicating a neuronal maturity level comparable to MNs derived in studies by Wang *et al*. and AnswerALS. In contrast, lower *RBFOX3* levels in MNs from Kiskinis *et al*. and Dash *et al*. suggested reduced maturation in those models. Notably, *MAPT* expression increased by Day 20 compared with Day 10, indicating enhanced maturity with prolonged culture.

We also assessed markers indicative of MNPs and MNs. The MNP markers *NKX6-1*, *NKX6-2*, which are involved in ventral neural tube patterning, *PAX6*, a regulator of neurogenesis and neural patterning, and *OLIG2*, essential for MN progenitor specification, were expressed at low levels, suggesting a limited population of MNPs. To evaluate MN identity, we analyzed the expression of *MNX1, ISL1* and *ISL2*, transcription factors critical for MN differentiation and survival, with *MNX1* serving as a key determinant of MN identity and maturation in early development; *NEFH*, a component for the neuronal cytoskeleton important for axonal integrity; *CHAT*, encodes a functional marker specific to cholinergic neurons, which include spinal MNs. Although *ISL1*, *ISL2*, and *CHAT* were expressed at lower levels, their presence, together with other neuronal markers, is consistent with successful MN differentiation.

Given that SOD1-related ALS primarily affects limb movements innervated by MNs in the lateral motor column (LMC), we further assessed markers associated with the LMC MN subtype, including *FOXP1*, *NTM*, *LHX1*, *HOXA5*, *RET*, *POU3F1*, and *ALDH1A2*. *FOXP1* is critical for LMC neuron formation (Adams et al. 2015), while *LHX1* distinguishes dorsal limb-innervating neurons. *NTM* facilitates axon guidance and limb innervation, and *ALDH1A2* is involved in early differentiation via retinoic acid synthesis, showed reduced expression, consistent with maturation. *POU3F1* (also known as *SCIP*) contributes to LMC MN maturation, increased over time, further supporting ongoing MN development. *RET* and *EPHA4*, critical for axon guidance, showed expected expression patterns, with *RET* serving as an early marker, and its reduction further supporting MN maturation. The expression levels of these markers in the differentiated MNs were relatively higher compared in previous studies of Kiskinis *et al*., Dash *et al*., and Wang *et al*., and were comparable to AnswerALS. The dynamic expression of *ALDH1A2*, *RET* and *POU3F1* further confirmed the progressive maturation of MNs.

We also evaluated the rostral cranial MN marker *PHOX2B* for comparison (Samad et al. 2004). Cranial nerve nuclei, located in the brainstem (Ragagnin et al. 2019), represent MN populations that are relatively less affected in *SOD1*-associated ALS. The result reveals a low representation of this subtype in the current samples, suggesting that the differentiation protocol predominantly favors spinal MN subtypes.

Since raw counts can be influenced by differences in study design, we reprocessed both our data and raw data from Dash, Wang, and Kiskinis et al. using the same alignment and quantification pipelines, and then normalized as Log_(TPM + 1) to facilitate accurate comparative analysis (Fig. 1H). To validate the identity of our differentiated MNs, we assessed potential contamination by non-MN subtypes through the expression of selected markers, including those for astrocytes and cholinergic, dopaminergic, GABAergic, glutamatergic, and serotonergic neurons. The results were consistent with those obtained from raw counts, showing detectable expression of astrocytic and various neuronal subtype markers across all datasets, indicating a proportion of non-MN cells. However, cholinergic neuron markers, such as *ACHE* and *SLC18A3*, were predominant, suggesting a MN-enriched population. Notably, markers for GABAergic (*GABBR1*) and glutamatergic (*GLUL*) neurons were also expressed at considerable levels.

Collectively, these findings confirmed the predominant identity and progressive maturation of the differentiated cells as MNs. Notably, marker expression levels were comparable between MNs with and without *SOD1* mutations, likely reflecting the early stage of disease pathology. This aligns with observations in ALS patients, who often retain normal motor function during the preclinical phase with minimal MN loss (Brown and Al-Chalabi 2017). Although some expression of non-MN markers (e.g., astrocytic, GABAergic, and glutamatergic) was detected, such heterogeneity is consistent with previous iPSC-based MN differentiation protocols, as demonstrated by our cross-dataset comparison, and is generally considered acceptable for downstream analyses. Thus, this model provides a validated platform for investigating the early impact of *SOD1* monogenic mutations in ALS.

### Transcriptomic Alteration of MNs with ALS-related *SOD1* mutations

Although few cellular pathological phenotypes were observed in our MN cultures, transcriptomic alterations may still be present. These early changes might not directly trigger neurodegeneration but could exacerbate cellular vulnerability and contribute to disease progression, complicating the understanding of *SOD1*-related mechanisms in ALS. To explore these alterations, we performed bulk RNA-seq to profile transcriptomic changes associated with *SOD1* mutations by comparing pooled data from WT/WT cells and from WT/D91A and WT/G94A mutants across days 10 and 20.

We found an anti-conservative distribution of p-values for DEGs, demonstrating distinct transcriptomic profiles between MNs with ALS-related *SOD1* mutations and controls (Supplemental Fig 2A). Principal component analysis (PCA) revealed that while the three groups with different SOD1 variants were distinct, the WT/WT samples clustered more closely with WT/D91A than with WT/G94A (Fig. 2A), indicating a greater transcriptional divergence associated with the G94A mutation. Volcano plot analysis identified 1,268 upregulated and 1,118 downregulated DEGs (adjusted p < 0.05), reflecting widespread transcriptional changes upon SOD1 mutation (Fig. 2B). RT-qPCR of selected genes showed a strong correlation with RNA-seq (Pearson R = 0.84 for Log_2_FC, p-value = 3.2 * 10^-11^) (Fig. 2C, Supplemental Fig 2B), confirming the accuracy of the RNA-seq data. Interestingly, analysis of the protein-protein interaction (PPI) network constructed from DEGs revealed no correlation between Log_2_FC and network connectivity (Fig. 2D). Notably, central hub genes such as *MYC*, *JUN,* and *SRC* exhibited only modest changes in expression, whereas the most dysregulated genes, including *GRIN3A*, *ALG10B*, and *PCDHGB5*, had relatively low connectivity within the network (Supplemental Fig 2C).

**Figure 2.**
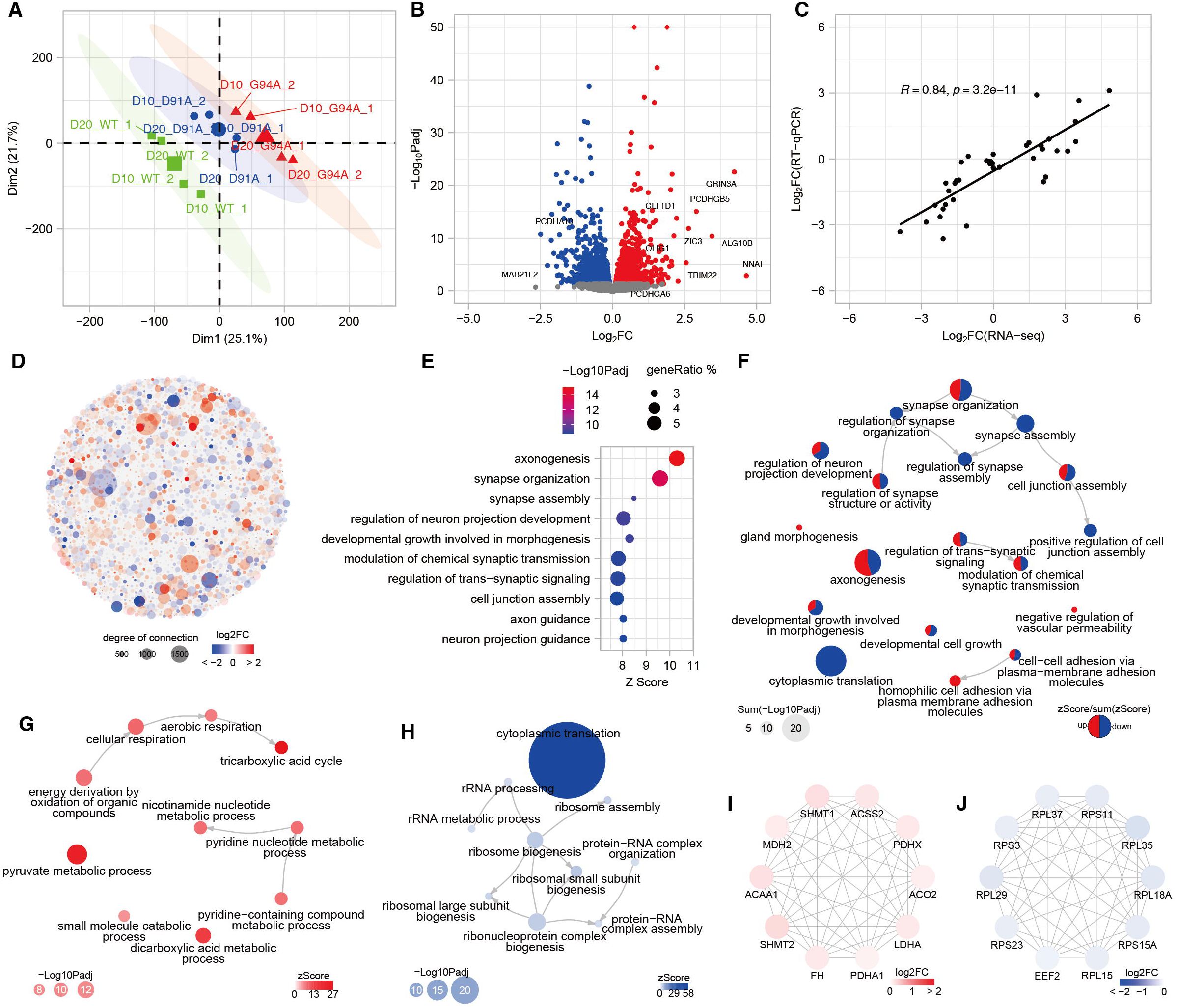
Pooled DEG analysis between MNs carrying mutated SOD1 and WT/WTSOD1. (**A**) PCA analysis of expression levels in MNs with different SOD1 variants. (**B**) Volcano plot showing genes differentially expressed between MNs with and without SOD1 mutations. (**C**) Pearson correlation between Log_2_FC from RNA-seq and RT-qPCR results for genes in Day 10 and Day 20 MNs, showing strong correlation. (**D**) PPI network for DEGs. Each node represents a gene. Node size indicates its degree (number of first neighbors), and color reflects Log_FC (mutant vs. WT/WT), capped at ±2 for visualization. (**E**) GO enrichment analysis for biological processes of DEGs between MNs with SOD1 mutations and WT. (**F**) Integrated directed graph of top enriched terms for both upregulated and downregulated gene set. Node size reflects combined −Log___Padj and color proportions indicate relative z-score contributions of each set. (**G**) The most significant module of STRING network for upregulated genes. (**G**) Directed graph for top enriched terms for upregulated genes. (**H**) Directed graph for top enriched terms for downregulated genes. (**I**) PPI network for the most connected genes in module up1. Node color reflects Log2FC. (**J**) PPI network for the most connected genes in module dn1.

To investigate the functional implications of gene expression changes, we performed GO enrichment analysis. DEGs were predominantly enriched in neuronal pathways, with nine of the top ten GOBP terms, ranked by −Log_10_(Padj), associated with neuronal processes, such as axonogenesis, synapse organization, and modulation of chemical synaptic transmission (Fig. 2E). GOCC analysis further revealed significant changes in neural structures including synapses and axons (Supplemental Fig S2D). To assess the directionality of these changes, we performed GOBP enrichment analysis separately for upregulated and downregulated genes. The term cytoplasmic translation was highly enriched in the downregulated genes (adjusted p = 8.27 * 10_¹_). Several pathways appeared among the top ten significantly enriched in both gene sets (Supplemental Fig S2E). To further explore the relationships among enriched biological processes, we constructed a directed graph illustrating the first-level parent-child relationships among the top enriched terms for both gene sets. The resulting network revealed a highly interconnected network characteristic, including synapse assembly, synaptic signaling, axonogenesis, and cytoplasmic translation (Fig. 2F). The hierarchical organization of these enriched GO terms suggested a coordinated alteration of synaptic structure and function, indicating that *SOD1* mutations may impair local protein synthesis and synaptic integrity in MNs.

Although the number of upregulated genes exceeded that of downregulated genes, the top enriched terms in the upregulated gene set were less significant, suggesting potential differences in the modular organization of PPI networks between the two sets. To further explore these network features, we applied molecular complex detection (MCODE) to identify densely connected modules within the PPI networks. Five highly interconnected modules (MCODE score ≥ 5) were identified among downregulated genes, and four among upregulated genes (Table 1). Notably, the downregulated genes formed denser modules, with the top module (dn1) exhibiting a high score of 42.591 and comprising 45 genes. In contrast, upregulated modules were smaller and less cohesive, suggesting a more functionally diverse or diffuse biological response.

**Table 1.**
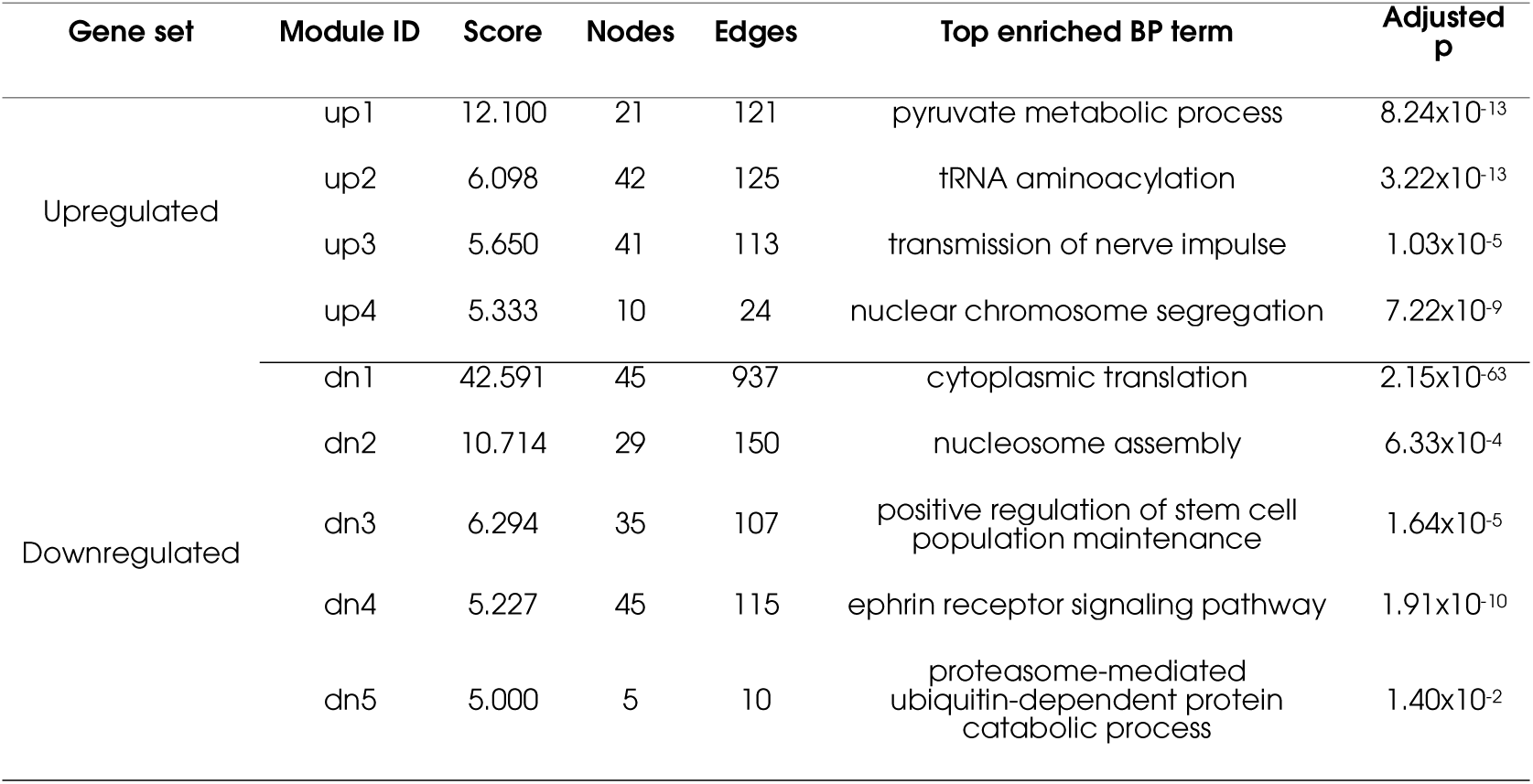
MCODE-Detected Modules in PPI Networks of Up- and Downregulated DEGs.

To characterize the functional indications of these clusters, we performed GOBP enrichment analysis. The most significant module for upregulated set was primarily enriched in energy production processes, including organic compound oxidation (Supplemental Fig 2F) with looser connected GO terms (Fig. 2G). Conversely, the top modules for downregulated sets were strongly enriched in cytoplasmic translation (adjust p = 2.15 * 10^-63^), with tightly interconnected terms associated with ribosome biogenesis (Fig. 2H, Supplemental Fig 2G).

To find the hub genes for these core modules, we extracted top ten genes with the most degrees of connections. *SHMT1*, *ACSS2*, *PDHX, ACO2*, *LDHA, PDHA1*, *FH*, *SHMT2*, *ACAA1* and *MDH2* were identified among module up1 (Fig. 2I). These genes are central regulators of glycolysis, the tricarboxylic acid (TCA) cycle, and one-carbon metabolism, indicating a coordinated upregulation of energy production pathways. This network-level prominence aligns with GO enrichment results, where the up1 modules were significantly enriched in aerobic respiration, pyruvate metabolic process, and nicotinamide nucleotide metabolism. The presence of these metabolic hub genes suggests an adaptive or compensatory response to cellular stress in ALS-related MNs. In contrast, the core module among downregulated genes was densely interconnected and dominated by ribosomal proteins (Fig. 2J), supporting a coordinated repression of cytoplasmic translation and ribosome biogenesis. Notably, most of these hub genes exhibited only modest changes in expression.

Collectively, these findings reveal distinct network architectures between up- and downregulated genes in *SOD1* mutant MNs. Downregulated genes formed dense PPI modules enriched in cytoplasmic translation and ribosome biogenesis, dominated by ribosomal protein hubs (e.g., RPS, RPL), suggesting coordinated repression of protein synthesis. This tightly connected network may reflect core neuronal functions particularly susceptible in ALS. In contrast, upregulated genes formed smaller, less cohesive modules enriched in oxidative metabolism, with metabolic hubs pointing to a compensatory or stress-adaptive response. Despite modest expression changes (|Log_FC| < 1), the high connectivity of these hub genes underscores their potential regulatory significance. Together, these opposing network features suggest that *SOD1* mutations disrupt MN function through translational repression coupled with metabolic adaptation.

### Features of DEG Clusters According to Different Mutations and Time points

Our previous research demonstrated that different *SOD1* variants correlate with varying ALS severities, as reflected by distinct disease durations (Huang et al. 2024). Specifically, patients with the WT/D91A mutation have an average disease duration of 125.71 ± 80.26 months, while those with the WT/G94A mutation experience a significantly shorter duration of 37.50 ± 20.68 months. To investigate the mechanisms driving these differences, including the temporal shifts of different pathways, we performed a comparative analysis of DEGs between WT/D91A and WT/WT, and WT/G94A and WT/WT, at Days 10 and 20 separately.

WT/D91A showed a reduction in DEGs over time, from 779 upregulated and 692 downregulated genes at Day 10 to 339 and 228, respectively, at Day 20 (Fig. 3A). This attenuation suggested a potential transcriptomic compensation or stabilization, aligning with the slower disease progression observed clinically. In contrast, WT/G94A exhibited a consistently elevated and expanding DEG burden, from 1,257 upregulated and 1,305 downregulated genes at Day 10 to 1,891 and 1,487 at Day 20, indicating persistent dysregulation that may contribute to its more aggressive ALS phenotype.

**Figure 3.**
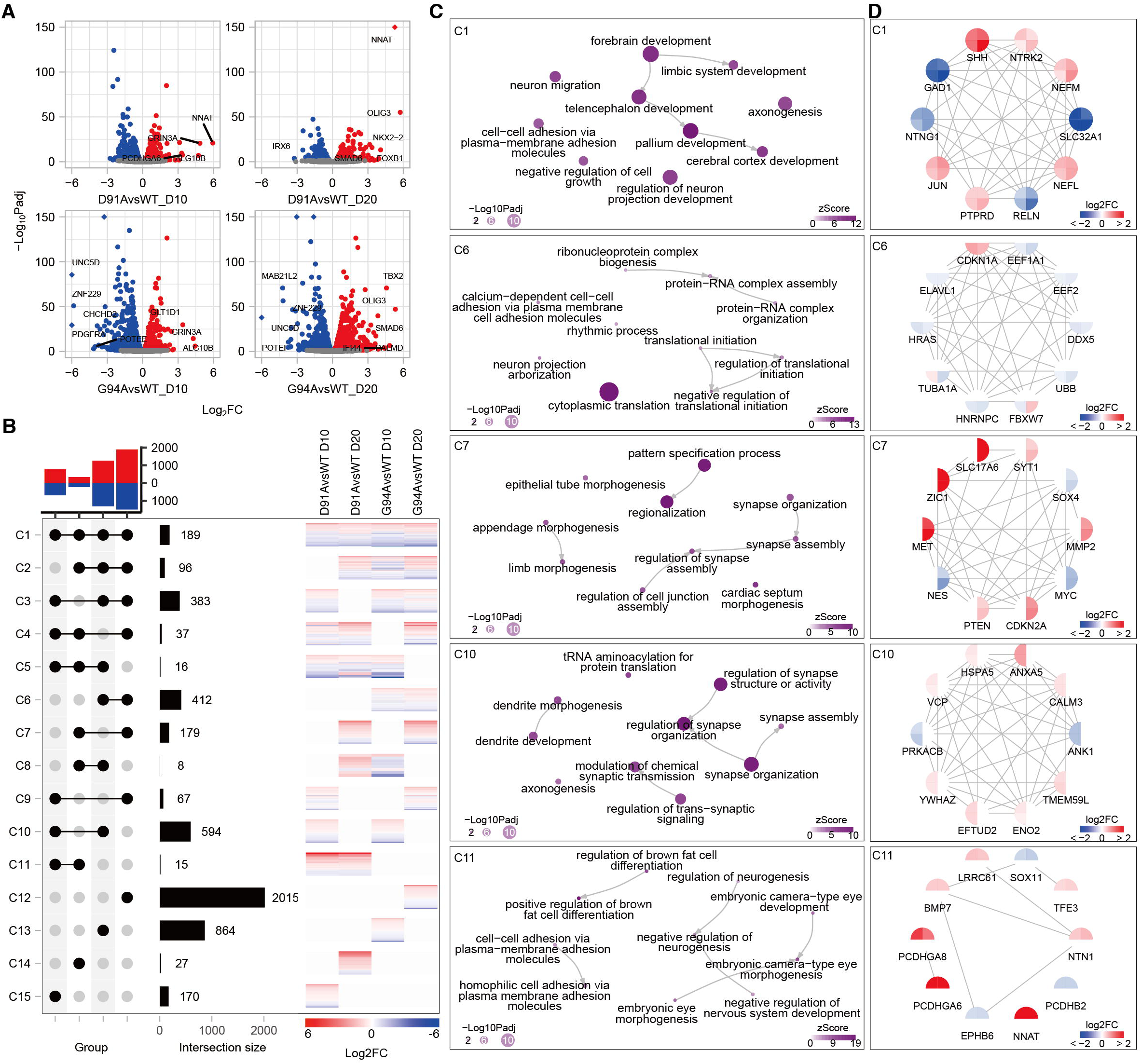
GO enrichment analysis of DEG clusters comparing SOD1 mutations at day 10 and 20. (**A**) Volcano plots of each comparison. (**B**) Upset plot showing overlap of DEGs across different comparison strategies. Heatmap of DEGs in each cluster, with the color bar representing Log_2_FC of expression level. (**C**) Directed graphs of GOBP parent-child relationships for selected gene clusters. (**D**) PPI networks for top ten connected genes within each cluster.

To explore the biological function of these changes, we clustered DEGs based on overlap across genotype-time comparisons (Fig. 3B), and representative clusters were analyzed for functional enrichment, and hub genes (Fig. 3C-D, Table 2). Cluster 1 (C1) captured 189 shared DEGs across both mutations and time points, enriched in forebrain and telencephalon development, axonogenesis, and cell adhesion pathways, reflecting the fundamental neurodevelopmental disruptions imposed by *SOD1* mutations. Hub genes in PPI network for this cluster showed consistent directional dysregulation across all conditions, upregulation of *SHH*, *NTRK2*, *NEFM*, *NEFL*, *JUN*, and *PTPRD*, and downregulation of *GAD1*, *NTNG1*, *SLC32A1*, and *RELN*. This consistent dysregulation suggests a shared transcriptional signature in ALS MNs, possibly reflecting subtype-specific vulnerability or a shift in neuronal identity involving neurodevelopmental, synaptic, and cytoskeletal pathways.

**Table 2.**
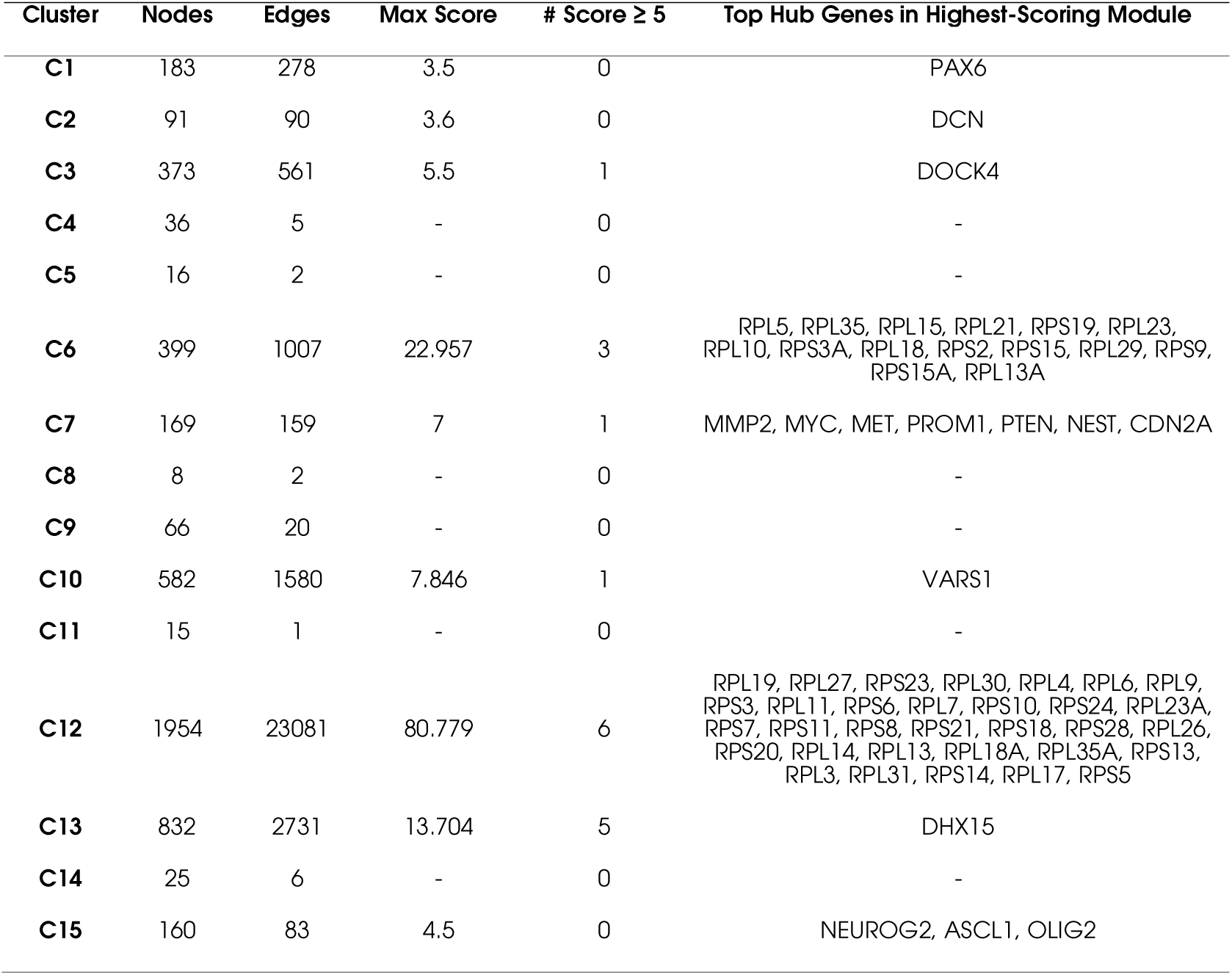
Network Characteristics of DEG Clusters.

Cluster 11 (C11) comprised 15 genes uniquely dysregulated in WT/D91A across both time points, enriched in neurogenesis and morphogenesis but lacking strong interconnectivity, suggesting subtle and disorganized transcriptomic changes aligned with the milder disease phenotype. The gene number was insufficient to construct a PPI network. The upregulation of NNAT, PCDHGA6, and PCDHGA8 suggested a localized activation of genes involved in neuronal development and cell adhesion. In contrast, Cluster 6 (C6), consisting of 412 genes specifically altered in WT/G94A, was enriched for cytoplasmic translation. Despite strong network connectivity, hub genes in C6 showed modest expression changes, indicating coordinated but less intense transcriptional shifts.

Temporal dynamics were further captured in Cluster 10 (C10) and Cluster 7 (C7). C10, comprising 594 genes dysregulated exclusively at Day 10 in both mutants, was enriched for early neurodevelopmental processes such as dendrite morphogenesis and synapse assembly. In contrast, C7 included 179 genes uniquely dysregulated at Day 20, associated with regionalization, pattern specification, and synaptic organization. Notably, despite fewer genes, C7 exhibited stronger dysregulation of hub genes than C10, potentially reflecting an intensification of pathogenic transcriptional programs during later stages of MN maturation. Together, these findings reveal both shared and divergent transcriptomic features across *SOD1* mutations. While both WT/D91A and WT/G94A perturb core neuronal development pathways, G94A induces more persistent and extensive dysregulation, especially in translational control, consistent with its more severe clinical manifestation. The presence of shared early alterations and distinct temporal trajectories underscores the importance of mutation-specific and time-sensitive mechanisms in ALS pathogenesis and suggests tailored intervention windows for therapeutic targeting.

### GSEA Reveal Key Role of Translation Inhibition in *SOD1*-related ALS Progression

Although GO enrichment of DEGs highlighted pathways affected by the WT/D91A and WT/G94A mutations across different time points of MN culture, such over-representation approaches are inherently biased toward genes with large expression changes. As noted earlier, hub genes within tightly connected networks often exhibited only modest expression changes. Consequently, conventional enrichment may miss biologically meaningful pathways that are subtly dysregulated. To gain a more comprehensive, system-level view of transcriptomic alterations, we performed gene set enrichment analysis (GSEA) using GSEApy across multiple databases, including GOBP, GOCC, GOMF, KEGG, Reactome, and Wikipathways and visualized the top two enriched terms from each comparison using a dot plot (Fig. 4A).

**Figure 4.**
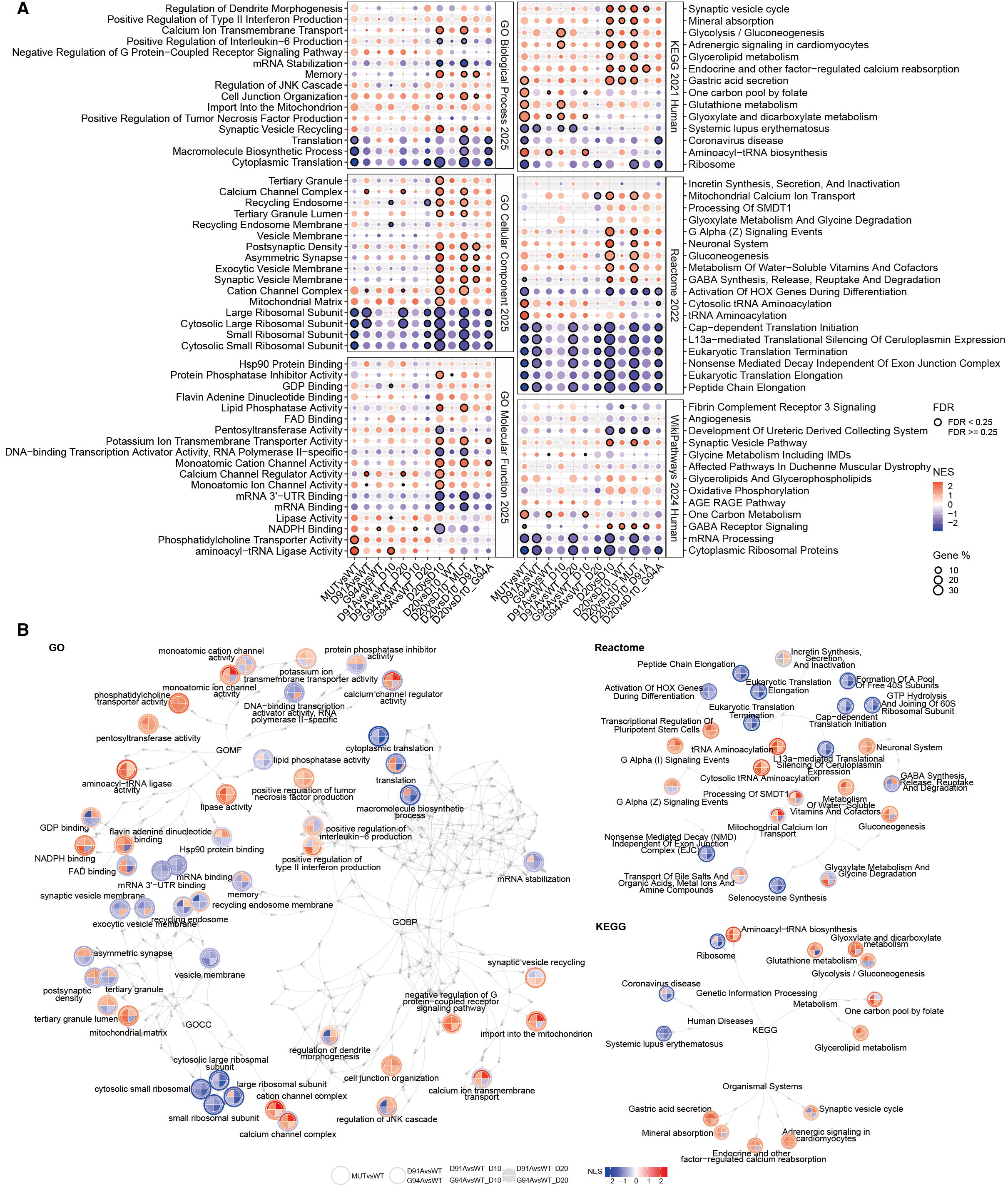
GSEA analysis reveals common downregulation of translation in SOD1 mutants. (**A**) Dot plot of GSEA analysis using databases including GOBP 2025, GOCC 2025, GOMF 2025, KEGG 2021, Reactome 2022 and WikiPathway 2024, top two terms for each comparison were displayed for every database. (**B**) Directed graph of the displayed terms in (A) and their ancestors. Edges indicate hierarchical relationships.

Most enriched terms exhibited stronger enrichment in the day 20 vs. day 10 comparisons, suggesting progressive transcriptomic remodeling. This trend was particularly evident in WT/D91A and WT/G94A compared to WT/WT. There were also different degrees of remodeling between the two mutations, for example, translation-related pathways were more strongly downregulated in WT/G94A (day 20 vs. day 10), whereas membrane-associated processes were more prominently upregulated in WT/D91A (day 20 vs. day 10).

To contextualize the enriched terms biologically, we mapped their hierarchical relationships using directed acyclic graphs, preserving all relevant ancestor terms to capture the broader functional context and lineage of each enriched category. (Fig. 4B). Among the databases, KEGG provided the most interpretable network, with enriched terms primarily related to genetic information processing, metabolism, organismal systems, and human diseases. Ribosome biogenesis was consistently downregulated, while aminoacyl-tRNA biosynthesis was upregulated in mutant MNs, suggesting a divergence in translation-associated processes. Several metabolic pathways, including glutathione, glycerolipid, glyoxylate and dicarboxylate metabolism, glycolysis/gluconeogenesis, and one-carbon pool by folate, were generally upregulated. Notably, glycolysis/gluconeogenesis was downregulated in WT/G94A but upregulated in WT/D91A; glutathione and glyoxylate/dicarboxylate metabolism were initially upregulated at day 10 but downregulated by day 20 in WT/G94A. Terms related to organismal systems, such as the synaptic vesicle cycle, were upregulated in mutants overall, while those linked to human diseases were downregulated.

GO networks further supported these findings. Translation-related terms, such as cytoplasmic translation, macromolecule biosynthesis, and mRNA stabilization (GOBP); cytosolic and ribosomal subunits (GOCC); and RNA 3’ UTR and mRNA binding (GOMF), were significantly downregulated, most prominently in WT/G94A at day 20 and least in WT/D91A at day 10. In contrast, aminoacyl-tRNA ligase activity (GOMF) was upregulated, though the degree declined over time and was stronger in WT/G94A than WT/D91A. Dysregulation of metabolism-related terms (e.g., mitochondrial import, matrix components, GDP/NADPH/FAD binding) was also evident. These terms were generally upregulated except in WT/G94A at day 20, where they were consistently downregulated. GDP binding was uniquely downregulated in WT/D91A at day 10 but upregulated at day 20, whereas it remained downregulated in WT/G94A. Ion transport and membrane trafficking were also affected, including altered expression of calcium/cation channel complexes and transporter activities (GOMF), as well as calcium ion transmembrane transport (GOBP). Membrane-related processes such as synaptic vesicle recycling, recycling endosomes, and exocytic vesicle membranes (GOCC) were dysregulated. Upregulated lipase activity and downregulated lipid phosphatase activity (GOMF) may contribute to these changes. Immune and cell communication pathways, including TNF, IL-6, IFN-γ production, GPCR signaling, cell junction organization, and the JNK cascade, were also impacted. Additional dysregulated terms included Hsp90 binding, RNA polymerase II-specific transcription activator activity, protein phosphatase inhibitor activity, and FAD binding. Notably, Hsp90 protein binding was downregulated in WT/G94A while upregulated in WT/D91A.

Reactome results corroborated these observations with finer resolution. Translation-associated terms, including peptide chain elongation, translation elongation and termination, 40S subunit pool formation, cap-dependent initiation, and L13a-mediated translational silencing, were downregulated in all comparisons, with more pronounced effects at day 20. Nonsense-mediated decay independent of exon junction complexes was similarly downregulated over time in both mutants. Conversely, tRNA aminoacylation pathways were upregulated, particularly at day 10. Metabolic pathways showed complex regulation. Water-soluble vitamin/cofactor metabolism and gluconeogenesis were upregulated, more so at day 10. Glyoxylate metabolism and glycine degradation were upregulated in day 10 but downregulated by day 20. Selenocysteine synthesis was downregulated in both, with a stronger effect at day 20. Neuronal system pathways were broadly upregulated, particularly at day 20, while GABAergic transmission was downregulated in WT/D91A at day 10 but later upregulated, contrasting with persistent downregulation in WT/G94A. Additional Reactome terms, such as mitochondrial calcium ion transport and SMDT1 processing, showed early upregulation in both mutants, further increasing in WT/D91A but decreasing in WT/G94A in day 20. Similar patterns were observed for bile salt and organic acid transport. GPCR signaling also showed subtype-specific effects: Gα(i) signaling was upregulated in both mutants, while Gα(z) signaling was upregulated in WT/D91A but downregulated in WT/G94A.

In summary, the SOD1 D91A and G94A mutations were associated with extensive transcriptomic alterations in MNs, including consistent downregulation of translation-related pathways and broader disruptions in metabolic, mitochondrial, and signaling processes. A notable feature was the progressive decline in the expression of genes involved in cytoplasmic translation by day 20, which may reflect reduced protein synthesis capacity and increasing cellular stress. Upregulation of tRNA aminoacylation pathways was observed at earlier stages, suggesting a potential compensatory response, although this upregulation appeared limited in duration and scope. Differences between D91A and G94A mutations were evident in the regulation of biosynthetic and metabolic processes, indicating mutation-specific patterns of cellular adaptation. These findings point to a dynamic interplay between adaptive and potentially maladaptive transcriptional responses that affect cellular homeostasis in a mutation- and time-dependent manner. While the functional consequences of these changes remain to be fully elucidated, the transcriptomic signatures observed here provide a foundation for understanding divergent molecular trajectories in SOD1-related ALS pathogenesis.

### Dysregulation of interactome for hubs and miRNAs

To explore upstream contributors to impaired cytosolic translation, we analyzed alternative splicing (AS) across five major event types using rMATS (Table 3). In D91A MNs at day 10, skipped exon (SE) events dominated with 1,596 gains and 1,248 losses, alongside moderate dysregulation in retained introns (RI), mutually exclusive exons (MXE), and alternative splice sites (A3SS/A5SS). At day 20, SE changes remained high (+1,167, −1,316), with increased disruption in MXE and A3SS. G94A mutants showed more pronounced AS alterations, particularly at day 10 (+2,133 SE, −1,273) and day 20 (+1,785 SE, −1,741; +403 MXE, −394). Notably, both mutants exhibited progressive RI accumulation, with increased gains and reduced losses over time. These data reveal extensive, dynamic splicing abnormalities in SOD1 mutant MNs, more severe in WT/G94A, consistent with its clinical phenotype.

**Table 3.**
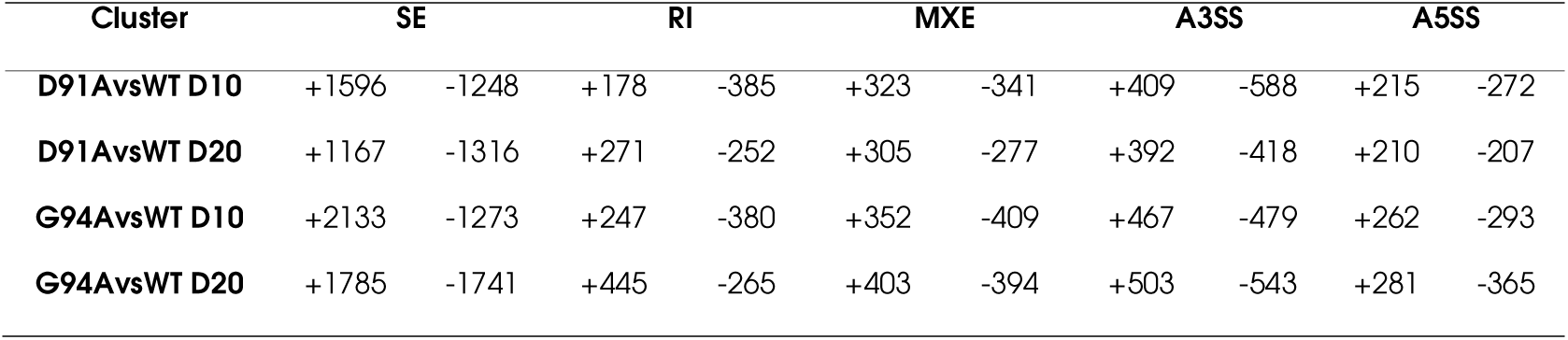
Alternative splicing events.

To further elucidate regulatory components, we conducted GSEA using the PPI_hub database to identify central network regulators. AGO1 and AGO2 emerged as significantly enriched hubs; both are core components of the RNA-induced silencing complex, mediating miRNA-guided post-transcriptional repression (Cheloufi et al. 2010). Given their centrality and known role in ALS-related gene regulation, we also examined miRNA regulatory dynamics across conditions using GSEA (Fig. 5A). Consistent with pathway-level GSEA, enrichment scores were generally stronger in day 20 vs. day 10 comparisons.

**Figure 5.**
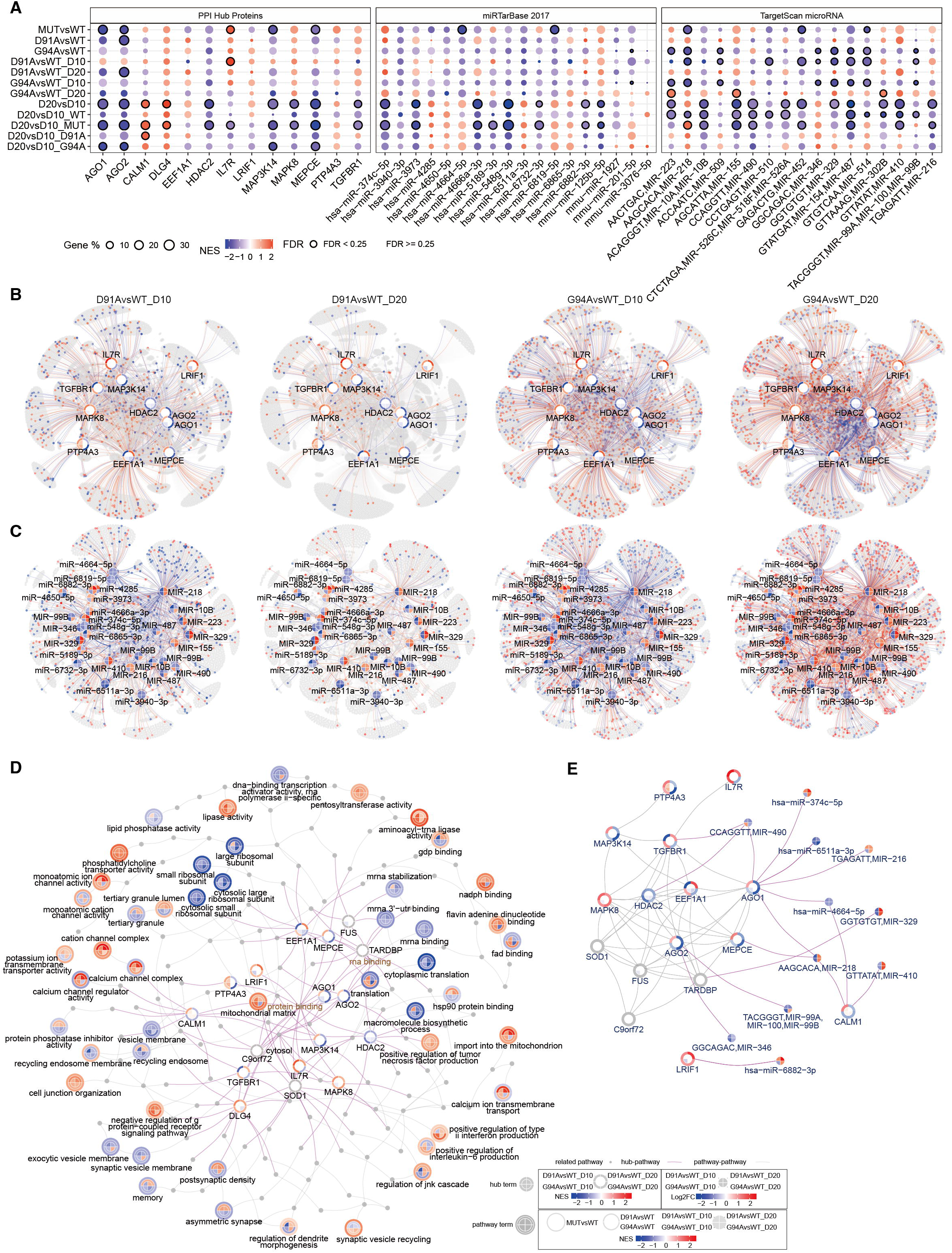
Alternative splicing analysis, hub gene interactome and miRNA target network analysis. (**A**) Dot plot of GSEA analysis using databases including PPI Hub Proteins, miRTarBase 2017 and TargetScan microRNA. (**B**) PPI network for enriched hub genes and their interactomes. Red for upregulated genes and blue for downregulated genes. (**C**) Network for enrich miRNAs and their targets. Red for upregulated genes and blue for downregulated genes. (**D**) Integrated undirected network for the hub-GO and GO subnetwork. (**G**) Integrated network for the hub and miRNA terms.

To validate GSEA findings, we constructed and visualized PPI networks for enriched hubs and miRNA-target networks (Fig. 5B-C). Most hub genes themselves showed mild expression changes. The PPI and miRNA regulatory networks recapitulated GSEA results, reflecting similar up- and down-regulated interactomes. Notably, the dysregulation of hub interactome declined from day 10 to day 20 in WT/D91A but increased in WT/G94A. A similar pattern was observed in the miRNA targets, which was predominantly downregulated at day 10 in both mutants but shifted toward upregulation at day 20, especially in WT/G94A.

To further elucidate the functional relevance of the enriched hub genes in ALS pathology, we integrated the gene-term network, incorporating both GSEA-identified hub genes and established ALS-associated genes (*SOD1*, *C9orf72*, *TARDBP*, and *FUS*), based on GO annotations with the undirected GO term-term network (Fig. 5D). A substantial proportion of these genes were annotated with RNA- and protein-binding functions, consistent with roles in post-transcriptional regulation. Translational processes were associated with FUS and TARDBP, both implicated in mRNA 3’-UTR binding; AGO2; and EEF1A1, a core translation elongation factor broadly connected to multiple translation-related GO terms. Beyond translation, SOD1 and CALM1 were annotated with mitochondrial matrix functions, with CALM1 also linked to calcium channel complexes, suggesting a role in calcium signaling and homeostasis. DLG4 was associated with the postsynaptic density, implicating synaptic dysfunction, while HDAC2 was linked to the positive regulation of tumor necrosis factor production, indicating potential involvement in inflammatory signaling.

To integrate the regulatory relationships between the identified hubs and miRNAs, we constructed a composite network combining PPIs among hub and ALS-associated genes with miRNA-target interactions (Fig. 5E). The PPI analysis revealed a densely interconnected network, suggesting that many of these hubs and ALS-associated genes function in concert or within shared regulatory modules. Overlaying miRNA-target data onto this framework further refined our understanding of gene regulation in ALS.

Among the miRNA-regulated nodes, AGO1 emerged as a central regulatory target, modulated by multiple miRNAs, including MIR-490, MIR-216, MIR-218, hsa-miR-374c-5p, hsa-miR-6511-3p, and hsa-miR-4664-5p. This highlighted AGO1 as a potential convergence point for miRNA-mediated regulation. Although AGO1 expression was only slightly downregulated in WT/G94A at day 10, the reduction coincided with the downregulation of targets associated with most of these miRNAs in the same condition. AGO1 expression remained relatively stable in other comparisons. CALM1 shared two miRNAs with AGO1, MIR-218 and hsa-miR-4664-5p, and was additionally targeted by MIR-410, implying coordinated post-transcriptional control of genes involved in calcium signaling and mitochondrial homeostasis. CALM1 was modestly upregulated at day 10 in both mutant lines, with a further increase in expression at day 20 in WT/G94A, consistent with miRNA-associated activity. MIR-218 also targeted MEPCE. MIR-490 regulated AGO1, TGFBR1, and HDAC2, indicating its broader involvement in RNA silencing, TGF-β signaling, and inflammatory pathways.

Other notable interactions included MIR-329 targeting TARDBP, MIR-346 and MIR-99A targeting AGO2, and hsa-miR-6882-3p targeting LRIF1. LRIF1 expression was downregulated in WT/G94A at day 10, consistent with the downregulation of hsa-miR-6882-3p targets. In contrast, LRIF1 remained stable in other conditions despite an overall elevation in hsa-miR-6882-3p targets, suggesting that miRNA activity may be modulated by context-dependent regulation or buffering mechanisms.

Collectively, our integrative network analyses demonstrate that translational repression in SOD1-linked ALS arises from converging disruptions in splicing, miRNA regulation, and hub gene interactions. Mutation-specific splicing defects and dynamic miRNA activity, particularly involving AGO1 and AGO2, reflect impaired post-transcriptional control. The combined PPI and miRNA networks reveal coordinated dysregulation across RNA processing, mitochondrial function, synaptic signaling, and inflammation. This complex regulatory landscape connects canonical ALS genes with central hubs and miRNAs, supporting a model in which shared circuits drive disease progression. These findings highlight key molecular nodes for therapeutic targeting and emphasize the value of GO-based and network-level approaches in elucidating ALS pathogenesis.

## Discussion

Despite the mixed nature of iPSC-derived MN cultures, isogenic iPSC lines offer a robust human in vitro model for studying early ALS pathogenesis with temporal resolution, an advantage over post-mortem tissues and animal models. Using precise genome editing, we generated iPSC lines with defined *SOD1* genotypes (WT/WT, WT/D91A, WT/G94A), establishing a genetically uniform platform for dissecting mutation-specific phenotypes without confounding variability. Comparison with published data confirmed that our MN cultures were acceptable for transcriptomic analysis. Future single-cell RNA-seq could further enhance cellular resolution.

Our comprehensive transcriptional analysis revealed that, despite the absence of overt cellular pathology in culture, *SOD1* mutations triggered complex, coordinated, and compensatory responses underlying the dysregulation of PPI and miRNA-gene regulatory networks. Notably, the most strongly dysregulated genes may not necessarily represent the principal drivers of pathology. Instead, subtle perturbations in hub genes could either reflect downstream consequences or act as potential initiating factors in disease mechanisms. We identified downregulated cytoplasmic translation in mutant MNs, consistent with findings in other ALS subtypes, including *C9orf72* (Goodman et al. 2019), *FUS* (Di Salvio et al. 2015), and *TARDBP* (Dash et al. 2022), indicating impaired protein synthesis as a common vulnerability. Both the pooled and separated analysis revealed the predominant role in translation downregulation caused by *SOD1* mutations. Coordinately downregulation of multiple translation related pathways according to GSEA analysis further makes the conclusion solid. Notably, aminoacyl-tRNA biosynthesis was upregulated, more at day 10 than day 20, likely reflecting a compensatory response to translational deficits. Therapeutic strategies aimed at restoring protein synthesis, such as using ISRIB (Marlin et al. 2024) or related agents, may offer cross-subtype benefit, including for *SOD1*-associated ALS.

Despite widespread dysregulation of translational pathways, metabolism, ion transport, and membrane trafficking, likely reflecting compensatory responses to impaired cytoplasmic translation, began to fail by day 20, particularly in the WT/G94A. While such changes have been observed as presymptomatic features (Spalloni et al. 2006; Lobsiger et al. 2007; Massignan et al. 2007; Murakami et al. 2007; Niessen et al. 2007), their role as compensatory responses has been underappreciated. Notably, the downregulation of key buffering systems, including glutathione metabolism, glyoxylate/dicarboxylate metabolism, and NADPH binding, may explain the limited efficacy of current ALS therapies targeting oxidative stress (Hogg et al. 2018). Interventions that support these systems, such as NAD^+^ precursors (Obrador et al. 2021), may offer added benefit, especially for aggressive ALS subtypes.

The downregulation of cytoplasmic translation might result from coordinated regulatory disruptions, including mutation-specific splicing defects, miRNA-mediated repression, and dysregulation of translation-associated hub gene networks. Intron retention may reduce the availability of mature transcripts for translation and is an established hallmark for ALS (Luisier et al. 2018). Temporal shifts in miRNA activity, with early hyperactivity and later attenuation, suggest a biphasic response. This may initially enhance silencing to buffer stress, followed by de-repression to support broader gene expression as translation deficiency persists. Although hub gene expression changes were modest, significant alterations in their interactomes indicate that functional consequences can result from disrupted network dynamics rather than expression level alone. These regulatory failures converge with mitochondrial impairment and redox imbalance, particularly in WT/G94A at later stages, to compromise translational capacity and reinforce the central role of post-transcriptional instability in ALS pathogenesis.

The precise mechanisms underlying translation downregulation in *SOD1*-mutant MNs remain unclear. One potential contributor is stress granule (SG) dysregulation, as SGs sequester RNA and translation-related proteins in response to stress (Taylor et al. 2016). SG pathology has been implicated in ALS cases involving *TARDBP* (Liu-Yesucevitz et al. 2010), *FUS* (Murakami et al. 2015), and *C9orf72* (Zhang et al. 2018) mutations. Although the link to *SOD1* is less well established, translational repression observed in *SOD1*-mutant MNs may perturb SG dynamics, leading to persistent SG formation that sequesters essential translational machinery and impairs stress recovery. Further investigation is needed to determine whether mutant SOD1 promotes SG formation and whether SG modulation offers a therapeutic avenue in *SOD1*-associated ALS.

The differential impact of D91A and G94A on cellular metabolic processes may be attributed to differences in protein stability and aggregation propensity. However, the precise mechanisms remain unclear and warrant further investigation. Although SOD1 enzymatic activity is not strictly required for ALS pathogenesis, loss of function may act as a disease modifier (Saccon et al. 2013). Our previous review (Huang et al. 2024) collected enzymatic activities of different variants from multiple studies and found that G94A exhibits a more pronounced loss of function compared to the functionally intact D91A, which may partially explain the greater severity associated with the G94A mutation. Whether modulating pathways altered in the D91A model could mitigate phenotypes observed in G94A remains an open question that warrants further investigation.

Finally, while ALS is generally considered to have an oligogenic basis (van Blitterswijk et al. 2012; Renton et al. 2014), our study shows that even monogenic *SOD1* mutations (D91A and G94A) can cause substantial transcriptional dysregulation in MNs. The G94A variant, in particular, shows greater disruption across multiple cellular pathways, correlating with its more severe clinical presentation. This highlights the importance of understanding the distinct mechanisms by which different SOD1 variants drive ALS progression. Such knowledge could guide the development of mutation-specific therapeutic strategies.

In summary, although some technological limitations remain, our iPSC-derived MN model presents an advanced platform for dissecting the mutation-specific mechanisms underlying ALS and our analysis introduces a potentially insightful pipeline for their investigation. By integrating transcriptomic profiling with network-based analyses and capturing temporal dynamics, the model underscores the potential for targeting translation dysregulation as therapeutic strategies and rescuing metabolism dysfunction as supplementary therapies. Targeting these early disruptions could provide valuable strategies for slowing ALS progression. Future research should further explore these pathways to develop stage-specific interventions that could slow or halt ALS progression.

## Methods

### Human iPSC culture

WT/D91A was purchased from WiCell with original name of WC034i-SOD1-D90A (Chen et al. 2014). All iPSC lines were fed with mTeSR1 (STEMCELL, 85850) daily and passaged with 0.5 mM EDTA (Invitrogen, 15575020) every two to three Days when reached a confluency of around 70% onto Matrigel (Corning, 354277)-coated 6-well plates (Corning, 3516) or confocal dishes (NEST, 801002) for immunofluorescence experiments. The cells were maintained in an incubator (Thermo Fisher Scientific) at 37 ℃ with 5% CO_2_. Ethical approval was approved by Medical Ethics Committee of Guangzhou Institute of Biomedicine and Health (GIBH), Chinese Academy of Sciences, reference number GIBH-IRB2020.

### Genome editing of iPSC

#### Design of sgRNA and ssODN

sgRNA and single-stranded oligodeoxynucleotides (ssODN) for homology-directed repair (HDR) were designed using the Alt-R™ CRISPR HDR Design Tool (IDT). The S.P Hifi Cas9 v3 nuclease (IDT, 1081060), requiring an NGG PAM site, was used. To select the optimal sgRNA for correcting D91A (D91A_WT), a 40 bp genomic sequence flanking the D91A (c.272A>C) site was analyzed, yielding four PAM sites on the positive strand and two on the negative strand. The sgRNA with the highest on-target score (80) and a TGG PAM was chosen, as also used by E. Deneault *et al*. (Deneault et al. 2021). This sgRNA was also applied to introduce G94A (c.281G>C), located 9 bp downstream, into the WT/D91A cell line. Two HDR templates (86 bp) were designed: one for D91A correction, and another for simultaneous correction of D91A and introduction of G94A with a K92K silent mutation (c.276A>G, Supplemental Table 1).

#### Preparation of RNP complex

The crRNA (IDT Alt-R® CRISPR-Cas9 crRNA XT), HDR donor oligo (IDT, Alt-R HDR Donor Oligo), and Cas9 electroporation enhancer (IDT, 1075915) were dissolved in IDT duplex buffer (IDT, 11-01-03-01) to 200 μM and vortexed. After brief centrifugation, the gRNA complex was prepared by mixing 2.5 μl of crRNA with 2.5 μl of tracrRNA, denaturing at 95°C for 5 minutes, and annealing at room temperature for 15-20 minutes. The RNP complex was then assembled by combining 2.25 μl of the annealed gRNA complex with 2 μl of Cas9 protein and incubating at room temperature for 10-20 minutes.

#### iPSC electro-transfection

iPSCs were prepared at 70-80% confluency in a 6-well plate for electro-transfection. Cells were treated with 10 μM Y27632 (STEMCELL, 72302) in mTeSR Plus (STEMCELL, 100-0276) three hours before the electro-transfection to enhance single cell viability post-digestion. A 24-well plate was coated with MatriGel for post-electro-transfection seeding. The seeding medium contained 1 μl Y27632 and 1.5 μl HDR Enhancer v2 (IDT, 10007910) per 1 ml mTeSR Plus. iPSCs were digested with 1 ml Accutase (STEMCELL 07920) at 37 °C for 5 minutes, followed by the addition of 5 ml pre-warmed DMEM/F12 (Gibco, 11330-032) to halt digestion. The cells were centrifuged, resuspended in DMEM/F12, and counted. One million cells were transferred to a new tube, centrifuged, and resuspended in Dulbecco’s Phosphate-Buffered Saline (DPBS), which was then removed. The P3 nucleofection buffer was prepared by mixing 82 μl Nucleofector solution with 18 μl supplement (Lonza, VPH-5012). Cells were resuspended in this buffer, and 2.5 μl of HDR template and 1 μl of Cas9 electroporation enhancer were added to the RNP complex. The 100 μl cell suspension was mixed with the RNP and transferred to a cuvette for electro-transfection using program B-016 on the Lonza Nucleofector 2b Device (Lonza AAB-1001). The mixture was seeded onto the MatriGel-coated well and incubated. The seeding medium was replaced with fresh mTeSR Plus 24 hours later and changed daily until reaching 70-80% density. Cells were then dissociated with EDTA and divided: one part for genome extraction, one for backup freezing in Stem Cellbanker (Amsbio, 11890), and one for subculturing in fresh mTeSR Plus for further experiments.

#### Preliminary verification of the editing efficiency

To preliminarily assess editing efficiency, genomic DNA was extracted using the Genomic DNA Micro Kit (Foregene, DM-01012). Cells were washed with DPBS and centrifuged at 300 g for 3 minutes, then digested with protease at 65°C for 30-60 minutes. The digest was transferred to a column, which was centrifuged to capture genomic DNA, followed by multiple washes to eliminate impurities. Finally, DNA was eluted with UltraPure™ DNase/RNase-Free Distilled Water (Invitrogen). Primers targeting exon 4 of the SOD1 gene were designed: forward primer Exon4_F (GTGTGTAGACGTGAAGCCTTG) and reverse primer Exon4_R (TCTGGATCTTTAGAAACCGCGA), yielding a 375 bp product. PCR was performed on the extracted genomic DNA from edited cell lines and the original WT/D91A reference using 2 × Taq Master Mix (Vazyme). The PCR products underwent Sanger sequencing at IGEbio (Guangzhou, China) to verify the mutations. Editing efficiency was evaluated using the Sanger sequencing results with the Indel Correction Efficiency (ICE) CRISPR Analysis Tool available at www.synthego.com.

#### Selection of monoclonal cell colonies

The middle 60 wells of three 96-well plates (Corning, 3598) were coated with MatriGel, while the edge wells were filled with DPBS to minimize evaporation. The subculture was dissociated into single cells using Accutase, and digestion was halted with DPBS. The cell mixture was centrifuged, resuspended in DPBS, and counted, then 180 cells were added to 36 ml of mTeSR Plus with 10 μM Y27632. This mixture was thoroughly mixed and dispensed into the precoated plates at 200 μl per well. Four Days post-seeding, the plates were examined under a microscope (Carl Zeiss, Axio Vert. 1). Wells with polyclonal colonies were excluded, while monoclonal colonies were marked and refreshed with mTeSR Plus daily. Once monoclonal colonies reached approximately 70% confluence (10-14 Days after seeding), half were mechanically dissociated and transferred to a freshly coated well, while the original wells were maintained and had their medium changed daily for 1-2 Days prior to lysis for genome extraction.

#### On-target sequencing

Genomic DNA was extracted and amplified from each monoclonal cell colony using the Animal Tissue Direct PCR Kit (Foregene, TP-01113). Colonies were washed with DPBS, lysed with proteinase in buffer AL at 65 °C, and the proteinase was deactivated at 95 °C. The lysate supernatant was then used for PCR with 2x PCR Easy Mix and primers Exon4_F and Exon4_R. PCR products were sent for Sanger sequencing to determine each colony’s genotype. Correctly edited colonies were expanded, frozen using programmed gradient cooling, and stored in liquid nitrogen for future studies.

#### Off-target sequencing

Off-target (OT) predictions were made using the online tool www.crispor.tefor.net (Haeussler et al. 2016), which employs cutting frequency determination (CFD) scores (Doench et al. 2016) based on the human genome (GRCh38) and a PAM sequence of 20bp-NGG for SpCas9, SpCas9-HF1, and eSpCas9 1.1. OT primers were designed using Primer3 on the same site, focusing on sites with CFD scores < 0.02, indicating low cleavage likelihood. The top 10 OT sites with suitable primers for a maximum amplicon length of 600 bp and a melting temperature of 60 °C were selected, including some exon regions despite low CFD scores, resulting in a total of 16 selected sites (Supplemental Table 2). To assess OT effects, PCR was conducted on the on-target colonies using the OT primers, with genomic DNA extracted via the Genomic DNA Micro Kit and amplified using 2 × Taq Master Mix. The PCR products were subsequently sent for Sanger sequencing.

#### Teratoma assay and Histological Analysis

iPSCs were dissociated with Accutase and resuspended at a concentration of 10 million cells per ml in mTeSR Plus medium, mixed with an equal volume of MatriGel. For intramuscular injection, NOD/SCID mice received 200 μl of the mixture at each injection site in the quadriceps muscle group. Mice were monitored weekly for teratoma size. Upon reaching approximately 1 cm in diameter, typically six weeks post-injection, mice were anesthetized with Tribromoethanol, and teratomas were surgically removed. The samples were measured, fixed in PBS with 4% paraformaldehyde (PFA, Beyotime, P0099), and embedded in paraffin. Sections were stained with hematoxylin and eosin (HE) for analysis. Paraffin section and HE staining were performed by Haozhao Liang in GIBH. This study was approved by the Laboratory Animal Welfare and Ethics Committee of GIBH (IACUC No. 2023046).

#### Karyotype analysis

Karyotype analysis was conducted by Tiancheng Zhou from Prof. Guangjin Pan’s lab at GIBH. iPSCs at 70-80% confluency were cultured in mTeSR1 with 0.2 μg/ml colchicine for 2 hours at 37°C to arrest cell division and promote chromosomal condensation. Cells were then dissociated with Accutase, fixed in 4% PFA, and placed on a glass slide. Chromosomes were spread, air-dried, and stained with Giemsa to enhance banding patterns. The chromosomes were subsequently examined, photographed, and paired to assess chromosomal intensity.

#### Western blot analysis

For protein expression analysis, iPSCs cells were lysed in RIPA buffer (Beyotime, B0013C) supplemented with protease and phosphatase inhibitors (ThermoFisher Scientific, 78429). Protein samples (20 μg) were separated by SDS-PAGE and transferred to a 0.45 μm PVDF membrane (MilliporeSigma, IPVH00010). The membrane was blocked with blocker solution (Bio-Rad, 1706404), followed by overnight incubation with primary antibodies against SOD1 (Invitrogen, PA5-27240) and GAPDH (CST, #2118) at 4°C. After washing with TBST, the membrane was incubated with secondary antibodies conjugated to horseradish peroxidase (HRP) at room temperature for 1 hour. Following the washing steps with TBST, the target protein bands were visualized using ECL Select Western Blotting Detection Reagent (Cytiva Life Sciences, 45-000-999) and Bio-Rad ChemiDoc Imaging Systems.

#### MN differentiation

MN differentiation was conducted according to the published protocol by Du *et al*.(Du et al. 2015) with a few modifications. The components of mediums are listed in Supplemental Table 3.

#### Neural induction

High-quality iPSCs were passaged one Day prior to induction (Day 0) using EDTA at a density of 10-20% and fed with mTeSR1. On differentiation Day (Day 1), mTeSR1 was replaced with medium #1, and the culture was washed with DMEM/F12 to remove dead cells if necessary. Medium #1 was changed daily for 5 Days, leading to the induction of neuroepithelial cells (NEPs) on Day 6. NEPs were dissociated with EDTA for 5 minutes, and after discarding the EDTA, cells were further detached by scraping in medium #2 due to their stronger adherence. Cells were then transferred to MatriGel-coated wells in a 1:3 to 1:6 ratio. Medium #2 was changed daily for 5 Days, differentiating NEPs into motor neuron progenitors (MNPs).

#### Expansion of MNP

MNPs were dissociated with prewarmed 0.5 mM EDTA for 5 minutes. EDTA was carefully discarded, and the cells were resuspended in medium #3. The resuspended cells were transferred onto MatriGel-precoated wells at a ratio of 1:3 to 1:6. MNPs could be expanded for at least five passages. Early passages of MNPs (P0, P1) were used for backup freezing, cells were resuspended in DMEM/F12 containing 10% fetal bovine serum (FBS, Gibco, 10270-106) and 10% dimethyl sulfoxide (DMSO, Sigma, D2650) and underwent programmed gradient cooling and stored in liquid nitrogen.

#### Maturation of MN

MNPs from passages 3-4 were used for differentiation. On Day six, MNPs were dissociated with EDTA and suspended in medium #4 before being transferred to uncoated wells at a 1:3 to 1:6 ratio. These cells spontaneously aggregated into spheres and were fed with medium #4, which was partially replaced daily for 6 Days. For subsequent passages, wells were coated overnight with 100 μg/ml poly-D-lysine (Sigma, P6407) and rinsed three times with double-distilled water. For immunofluorescence experiments, 96-well PhenoPlates (PerkinElmer, 6055302) were coated, while 6-well plates were prepared for protein and RNA extraction. After drying for 6 hours, the bottoms were coated with 10 μg/ml laminin (Sigma, L2020) for 4 hours.

On the Day of MN induction, cell spheres were transferred to a 15 ml centrifuge tube and centrifuged at 200 g for 3 minutes. The supernatant was discarded, and 1 ml of prewarmed Accutase with 50 μg/ml DNase I (Sigma, DN25) was added to the pellets. The mixture was pipetted to disperse the spheres and digested at 37 °C for 10 minutes. Digestion was stopped with DMEM/F12, and cells were centrifuged at 300 g for 3 minutes. After resuspension in medium #5, the cell suspension was filtered through a 40 μm strainer (Corning, 352340), counted, diluted, and seeded onto pre-coated plates at a density of 0.5-0.6 million cells per ml, with medium changes every three Days.

#### Immunofluorescence experiment

The culture medium was aspirated, and cells were fixed with 4% PFA for 15 minutes at room temperature. PFA was then removed and disposed of according to chemical waste guidelines, and cells were washed three times with DPBS for 10 minutes each. Cells were blocked for 1 hour at room temperature using a buffer of 1% BSA (Biosharp, BS114) and 0.1% Triton X-100 (Thermo Fisher Scientific, 85111) in DPBS. Cells were incubated with primary antibodies diluted in blocking buffer overnight at 4 °C. The primary antibodies used were OCT4 (Invitrogen, MA5-31458), SOX2 (Invitrogen, PA1-094), SSEA4 (Invitrogen, 41-4000), and Nanog (Invitrogen, PA1-097). After being washed with DPBS for 10 minutes three times, cells were incubated with secondary antibodies and 10 μM Hoechst 33342 (Thermo Fisher Scientific, H1399) in blocking buffer overnight at 4 °C. Samples were mounted with antifade medium (Beyotime, P0126) washed with DPBS for microscopy analysis using a LSM800 confocal microscope (Carl Zeiss) and a SpinSR super-resolution microscope (Olympus).

### Transcriptome analysis of MN

#### RNA-seq library construction and sequencing process

RNA extraction from MNs at Days 10 and 20 was performed following DPBS washes to remove mediums, with samples prepared in duplicate. Cells were lysed using RNA isolator solution (Vazyme, R401-01) on ice, scraped, and transferred to pre-cooled 1.5 ml centrifuge tubes. The lysate was flash-frozen in liquid nitrogen and transported to Guangzhou IGE Biotechnology Company for RNA-seq under dry CO_ conditions. RNA extraction and quality control were conducted using Nanodrop for concentration and purity assessment, and the Agilent 2100 Bioanalyzer for integrity evaluation. mRNA was enriched with capture beads, fragmented, and used as a template for synthesizing single-stranded cDNA, which was subsequently converted to double-stranded cDNA and purified.

Purified cDNA underwent end-repair, A-tailing, and junction connection, followed by size sorting with DNA clean beads. PCR amplification produced the final cDNA library, with preliminary concentration assessed via Qubit 3.0 and insert sizes analyzed with the Agilent 2100 Bioanalyzer. Libraries passing quality control were accurately quantified using the ABI Step One Plus real-time PCR system.

After quality assessments, libraries were pooled based on effective concentrations to achieve a target data volume of 6G for Illumina NovaSeq 6000, PE150 sequencing. This high-throughput paired-end sequencing enabled sequence determination at each end of the DNA fragments, facilitating assembly and comparative analyses. Raw image data from RNA-seq were processed through base calling analysis to generate raw Fastq data.

#### Real time quantitative PCR (RT-qPCR)

Total RNA was lysed using RNA isolator solution (Vazyme, R401-01) on ice followed by extraction using chloroform and isopropanol. Reverse transcription was carried out with 1 μg of RNA using HiScript II Q RT SuperMix for qPCR (Vazyme, R222). RT-qPCR was performed with ChamQ SYBR qPCR Master Mix (Vazyme, Q311) on a Bio-Rad CFX Connect Real-Time PCR Detection System, following the protocol: an initial step at 95 °C for 5 min, then 40 cycles of 95 °C for 10 s and 60 °C for 40 s. Gene expression was normalized to GAPDH, with primer sequences listed in Supplemental Table 4.

#### RNA-seq data analysis

Illumina adapters and low-quality raw Fastq data were trimmed using Trim Galore v0.6.5 in paired mode, applying a quality threshold of 25. Qualified files were aligned to the hg38.ensembl v81 reference genome using STAR (Dobin et al. 2013) v2.7.9a for gene and transcript analysis. Expected gene counts were quantified using RSEM (Li and Dewey 2011) (v1.3.3). Data from GEO (Supplemental Table 5) were processed using Fastq files following the same workflow, while expected gene counts from AnswerALS were directly accessed (access date: Aug 10, 2023, Supplemental Table 6). The expected gene counts and TPM were integrated into expression tables in R v4.4.1, with batch effects mitigated using the removeBatchEffect function from the limma package (Ritchie et al. 2015) v3.60.4. Ensembl IDs were mapped to their corresponding symbols using the AnnotationDbi package 1.66.0 with the org.Hs.eg.db 3.19.1 database. Differential alternative splicing analysis were performed using rMATS-turbo v4.3.0 (Wang et al. 2024).

Differential expression analysis in RNA-seq was performed using DESeq2 (Love et al. 2014) v1.44.0 to compare samples with and without SOD1 mutations. To mitigate batch variation, the removeBatchEffect function from limma was applied. Histograms of p-value distributions from both analyses were generated. Principal component analysis (PCA) was visualized using the fviz_pca function from the factoextra package v1.0.7, and heatmaps were constructed with the pheatmap package v1.0.12. Volcano plots were created using ggplot2 v3.5.1, applying a false discovery rate (FDR) cutoff of 0.05 for differentially expressed genes (DEGs). DEGs were analyzed for enrichment in subsets of Gene ontology (Ashburner et al. 2000) (GO) including biological processes (BP), molecular functions (MF), cellular components (CC) using the clusterProfiler package (Wu et al. 2021) v4.12.6, with both p-value and adjusted p-value (Benjamini-Hochberg) cutoffs set at 0.05. GO term network analysis were performed using package GO.db v3.19.1 and visualized using igraph v2.1.2, pie plot was added using package plotrix 3.8-4. Upset were generated using ggplot2.

#### STRING analysis and Module Identification

STRING networks were constructed based on 9606.protein.links.v12.0.txt downloaded from string-db.org and visualized by igraph v2.1.2. The most significant modules were identified using the Cytoscape plugin MCODE (Bader and Hogue 2003).

#### Gene set enrichment analysis (GSEA)

GSEA was conducted using the signal-to-noise method in GSEApy v1.1.3 (Python v3.12.2), utilizing databases for the three GO domains (GO_Biological_Process_2025, GO_Cellular_Component_2025, GO_Molecular_Function_2025) and pathway databases such as Wikipathways_2024_Human, KEGG_2021_Human, and Reactome_2022. For hub protein and miRNA analysis, the PPI_Hub_Proteins, miRTarBase_2017, and TargetScan_microRNA databases were employed. The top two enriched gene sets were extracted from each comparison. Visualization was achieved with a ggplot2. Network analysis of miRNA and hub interactions was performed using data from the STRING and TargetScan databases (accessed on October 16, 2024) as well as using package multiMiR v2.4.0 and visualized with igraph in R.

## Data access

All raw and processed sequencing data generated in this study have been submitted to the NCBI Gene Expression Omnibus (GEO; https://www.ncbi.nlm.nih.gov/geo/) under the accession number GSE283132.

## Acknowledgements

We thank Dr. Qiuyuan Fang and Professor Wei Yang from Zhejiang University for their valuable suggestions on genome editing. We also extend our gratitude to Tiancheng Zhou in Professor Guangjin Pan’s lab and Haozhao Liang from GIBH for their assistance with the karyotyping and teratoma formation assays to qualify the iPSCs. Additionally, we appreciate Assistant Professor Mingrui Ding from School of Life Science and Technology, ShanghaiTech University for the insightful discussions on the RNA-seq analysis.

This work was supported by the Research Funds from Health@InnoHK Program launched by Innovation Technology Commission of the Hong Kong SAR, P. R. China, Macau Science and Technology Development Fund (0061/2021/A2), and Shenzhen Science and Technology Innovation Commission (EF026/ICMS-SHX/2022/SZSTIC).

## Supplementary material

Supplementary material is available at *Genome Research* online.

