## Supplemental Fig legend for "Comparative Temporal Transcriptomic Analysis of *SOD1* Mutations in iPSC-Motor Neurons"

**Supplementary Figure 1. Establishment of Isogenic iPSC lines carrying different SOD1 Variants.** (**A**) Diagram of strategies for genome editing. (**B**) ICE prediction of genome editing efficiency based on Sanger sequencing results of the SOD1 gene for the first passage of mixed cells using two different strategies. (**C-D**) Characterization of characterized cell lines, including Sanger sequencing, microscopy, karyotyping, and teratoma formation results. **C** for WT/WT, and **D** for WT/G94A. (**E**) Western blot analysis for WT/WT, WT/D91A, and WT/G94A cell lines. (**F-G**) Immunofluoresence analysis of iPSC markers using primary antibodies against OCT4, SOX2, SSEA4, and Nanog. (**H**) RNA-seq based expression (Log₂(Count + 1)) for various cell markers across datasets.

**Supplementary Figure 2. Confirmation of DEGs between MNs with mutated SOD1 and wild-type SOD1.** (**A**) P-value distribution of DEGs, showing an anti-conservative pattern. (**B**) Fold change of selected genes calculated using RNA-seq and RT-qPCR data, normalized to GAPDH. (**C**) Scatter plot of DEGs showing their degree of connectivity (node degree) versus absolute Log_2_FC. (**D**) GOCC enrichment of DEGs. (**E**) GOBP enrichment for the upregulated and downregulated gene sets. (**F**) GOBP enrichment for the most significant upregulated cluster. (**G**) GOBP enrichment for the most significant downregulated cluster.
