## Supplemental Table for "Comparative Temporal Transcriptomic Analysis of *SOD1* Mutations in iPSC-Motor Neurons"

**Supplemental Table 1 Sequence for sgRNA and ssODN**

| sgRNA | Strategy | HDR Template |
| --- | --- | --- |
| AATGTGACTGCTGCCAAAGA | D91A_WT | TTAGGCATGTTGGAGACTTGGGCAATGTGACTGCTG**A**CAAAGATGGTGTGGCCGATGTGTCTATTGAAGATTCTGTGATCTCACTC |
|  | D91A_G94A | TTAGGCATGTTGGAGACTTGGGCAATGTGACTGCTG**A**CAA**G**GATG**C**TGTGGCCGATGTGTCTATTGAAGATTCTGTGATCTCACTC |

**Supplemental Table 2 Primers for OT sites**

| Site | Gene | Region | N* | CFD | Primer | Primer Sequence | Tm |
| --- | --- | --- | --- | --- | --- | --- | --- |
| chr10_100581074 | HIF1AN\|Metazoa_SRP | Intergenic | 3 | 0.69 | OT01_F | TCGTCGGCAGCGTCGTGGAGGTGGAGAGCGTAAA | 59.3 |
|  |  |  |  |  | OT01_R | GTCTCGTGGGCTCGGCCTGGTTGGAGGTCAAAGGT | 59.5 |
| chr4_166514221 | RP11-217C7.1 | Intron | 4 | 0.51 | OT02_F | TCGTCGGCAGCGTCTGACCAGAGTTCAGGCTTGG | 59.6 |
|  |  |  |  |  | OT02_R | GTCTCGTGGGCTCGGTCTCAAGAGTCCGCTCCTGT | 60.2 |
| chr10_67385057 | CTNNA3\|MIR7151 | Intergenic | 3 | 0.42 | OT03_F | TCGTCGGCAGCGTCTGTGCAGGCTCAAAGTTCCA | 60.1 |
|  |  |  |  |  | OT03_R | GTCTCGTGGGCTCGGAGCAAGCCATCTCTTTCCCT | 59 |
| chr10_67385158 | CTNNA3\|MIR7151 | Intergenic | 3 | 0.42 | OT04_F | TCGTCGGCAGCGTCCTGAGCCAGTGAAGAAAGGGA | 59.6 |
|  |  |  |  |  | OT04_R | GTCTCGTGGGCTCGGGGCCCTGGGATGCCATTATT | 60.1 |
| chr5_59270231 | PDE4D | Intron | 4 | 0.41 | OT05_F | TCGTCGGCAGCGTCAATCCATTGACCTGCCTTGC | 58.8 |
|  |  |  |  |  | OT05_R | GTCTCGTGGGCTCGGTCTGCAAGCTCTGTAAGGCT | 59 |
| chr5_89709956 | RP11-116A1.1\|AC112179.1 | Intergenic | 2 | 0.4 | OT06_F | TCGTCGGCAGCGTCAATGAAGTTCCCTGGGCCTC | 59.6 |
|  |  |  |  |  | OT06_R | GTCTCGTGGGCTCGGCCCTAAGCCCAAAGACACCA | 59.5 |
| chr12_63995693 | SRGAP1 | Intron | 4 | 0.39 | OT07_F | TCGTCGGCAGCGTCTGTCACCCAATGCATAGGGT | 59 |
|  |  |  |  |  | OT07_R | GTCTCGTGGGCTCGGAGGTCAGAGCCTTAGAGATCTGT | 60 |
| chr16_70077003 | PDXDC2P\|PDPR | Intergenic | 4 | 0.35 | OT08_F | TCGTCGGCAGCGTCTGAGAATGGCCCATCTGGTG | 59.7 |
|  |  |  |  |  | OT08_R | GTCTCGTGGGCTCGGTGAGAGGGATTCCAGCACTT | 58.3 |
| chr16_74554857 | GLG1 | Intron | 4 | 0.35 | OT09_F | TCGTCGGCAGCGTCTGAGAGGGATTCCAGCACTT | 58.3 |
|  |  |  |  |  | OT09_R | GTCTCGTGGGCTCGGTGAGAATGGCCCATCTGGTG | 59.7 |
| chr12_41555706 | PDZRN4 | Exon | 4 | 0.34 | OT10_F | TCGTCGGCAGCGTCGGGTCCAGAGAGTCCTAGCA | 60 |
|  |  |  |  |  | OT10_R | GTCTCGTGGGCTCGGTTGGACAGATCGTGGGGAAC | 59.6 |
| chr1_43447138 | SZT2 | Exon | 3 | 0.26 | OT11_F | TCGTCGGCAGCGTCAGGGAGGCTAGAATGGGGAG | 60.1 |
|  |  |  |  |  | OT11_R | GTCTCGTGGGCTCGGGGACAGAAACATGGGGACGT | 59.9 |
| chrX_24996791 | POLA1 | Exon | 4 | 0.14 | OT12_F | TCGTCGGCAGCGTCCCAGGAGTGGCTGTCCTTTT | 59.8 |
|  |  |  |  |  | OT12_R | GTCTCGTGGGCTCGGTCACCACAACTGCTTTCCCT | 59.4 |
| chr6_40407855 | RNU6-250P | Exon | 3 | 0.1 | OT13_F | TCGTCGGCAGCGTCAGCACCAGCCTCCTTGAAAA | 59.8 |
|  |  |  |  |  | OT13_R | GTCTCGTGGGCTCGGGACCACCCAAGCCCATAGAC | 60.1 |
| chr19_31278046 | TSHZ3 | Exon | 4 | 0.05 | OT14_F | TCGTCGGCAGCGTCCCGGCTCCTTCATCTTCTCC | 59.8 |
|  |  |  |  |  | OT14_R | GTCTCGTGGGCTCGGCCATCACAACCCTGCTGGAT | 60 |
| chr6_29355673 | OR5V1 | Exon | 4 | 0.02 | OT15_F | TCGTCGGCAGCGTCCAGGTGGGAGGCACATGTAG | 60.1 |
|  |  |  |  |  | OT15_R | GTCTCGTGGGCTCGGGTGACTGATCCACACCTGCA | 59.9 |
| chr17_6464721 | PITPNM3 | Exon | 4 | 0.01 | OT16_F | TCGTCGGCAGCGTCTCTGTGTCCAGGTGTACCCA | 60.1 |
|  |  |  |  |  | OT16_R | GTCTCGTGGGCTCGGCCTTCCAGAATGTCACGGCT | 60 |

*N = number of mismatched sites

**Supplemental Table 3 Mediums for MN differentiation**

| Reagent | Item No. | #1 | #2 | #3 | #4 | #5 |
| --- | --- | --- | --- | --- | --- | --- |
| DMEM/F12 | Gibco, 11330-032 | 24 ml | 24 ml | 24 ml | 24 ml | 24 ml |
| NeuroBasal | Gibco, 21103049 | 24 ml | 24 ml | 24 ml | 24 ml | 24 ml |
| N2 | Gibco, 17502-048 | 250 μl | 250 μl | 250 μl | 250 μl | 250 μl |
| B27 | Gibco, 17504-044 | 500 μl | 500 μl | 500 μl | 500 μl | 500 μl |
| Glutamax I | Gibco, 35050061 | 500 μl | 500 μl | 500 μl | 500 μl | 500 μl |
| L-Ascorbic acid | Sigma, A4544 | 0.1 mM | 0.1 mM | 0.1 mM | 0.1 mM | 0.1 mM |
| CHIR99021 | Tocris, 4423 | 3 μM | 1 μM | 3 μM |  |  |
| DMH1 | Tocris, 4126/10 | 2 μM | 2 μM | 2 μM |  |  |
| SB431542 | Stemgent, 04-0010 | 2 μM | 2 μM | 2 μM |  |  |
| RA | Tocris, 04-0021 |  | 0.1 μM | 0.1 μM | 0.5 μM | 0.5 μM |
| Purmorphamine | Stemgent, 04-0009 |  | 0.5 μM | 0.5 μM | 0.1 μM | 0.1 μM |
| VPA | Stemgent, 04-0007 |  |  | 0.5 mM |  |  |
| Coumpound E | Tocris, 6476 |  |  |  |  | 0.1 μM |

**Supplemental Table 4 Primers for RT-qPCR**

| **Target** | **Forward Primer sequence** | **Reverse Primer sequence** |
| --- | --- | --- |
| UNC5D | GCTCGACTCTAAGAACTGCACAG | TGAGGATTGCCACGACCACGAA |
| PCDH10 | TCTCCAACGGAAGCATTTTGTCC | CTATGTCGGCTTCCTGGAATGC |
| MAB21L2 | CGAAGGGTTCAACTTGCTCTCG | ACTGAGAGGCACTTGTTTCGGC |
| SLC32A1 | CTGGAACGTGACCAACGCCATC | TCATTCTCCTCGTACAGGCACG |
| SLC6A5 | CTGATGCTCCTCACTCTTGGAC | TGCGTAGGTACTTGGGAAACTCG |
| GRIN3A | ACACGGCAAACTTGGCTGCTGT | CTTCAGCACTGCTTTCTCGGAC |
| NNAT | GTGTTCCTGGAATGCTGCATTTAC | GACACCGTGTATGCCAGCTTCT |
| GLT1D1 | CTCATCTTGGCTGAGAACTGCG | TCAGTTCCACCAAAGATGACTCC |
| PCDHB5 | CCAGGCTGAAAACCGAGCACAA | GGCTGTTGTTCTCTCGGACGAA |
| PCDHA4 | CTACTCGTTGGTGCTGGACAGT | AGCGTTGTCGTTCACATCAGCC |
| ESRRB | GACATTGCCTCTGGCTACCACT | CTCCGTTTGGTGATCTCGCACT |
| RPL19 | TCACAGCCTGTACCTGAAGGTG | CGTGCTTCCTTGGTCTTAGACC |
| GAPDH | GTCTCCTCTGACTTCAACAGCG | ACCACCCTGTTGCTGTAGCCAA |

**Supplemental Table 5 Samples from GEO set**

| Sample | Donor | GEO set | Platform | Description | Ref |
| --- | --- | --- | --- | --- | --- |
| Kiskinis_A5V_1 | 39b-SOD1^+/A5V^ | GSE54409 | Illumina HiSeq 2000 (Homo sapiens) | Differentiated → plated on a glial monolayer → transduced with an Hb9::RFP lentiviral reporter → FACS after 15 days of additional culture | ^1^ |
| Kiskinis_A5V_2 | 39b-SOD1^+/A5V^ |  |  |  |  |
| Kiskinis_WT_1 | isogenic control |  |  |  |  |
| Kiskinis_WT_2 | isogenic control |  |  |  |  |
| Kiskinis_WT_3 | isogenic control |  |  |  |  |
| Wang_D91A_1 | ND29149-SOD1^+/D91A^ | GSE95089 | Illumina HiSeq 2500 (Homo sapiens) | DIV0-3: medium 1 →DIV3-6: RA added →DIV6: dorsomorphin removed, and SAG added→ DIV8: reseed on MatriGel-coated plates→ DIV9: DAPT and laminin added → DIV12: RNA isolation | ^2^ |
| Wang_D91A_2 | ND29149-SOD1^+/D91A^ |  |  |  |  |
| Wang_WT_1 | isogenic control |  |  |  |  |
| Wang_WT_2 | isogenic control |  |  |  |  |
| Dash_R116G_1 | SOD1 R115G_D8.9 | GSE210969 | Illumina HiSeq 2500 (Homo sapiens) | Total DIV30 | ^3^ |
| Dash_D91A_1 | SOD1 D90A_MM41 |  |  |  |  |
| Dash_CTRL_1 | Ctrl1_T122ctrl11 |  |  |  |  |
| Dash_CTRL_2 | Ctrl2_301ctrl21 |  |  |  |  |
| Dash_CTRL_3 | Ctrl3_4242ctrl31 |  |  |  |  |

**Supplemental Table 6 Samples from AnswerALS**

| Sample | GUID | Sex | Onset | Age |
| --- | --- | --- | --- | --- |
| Ans_A5V_1 | NEUCD502BFU | Male | 44 | 44 |
| Ans_G94A_2 | NEUCT842RJV | Male | 57 | 57 |
| Ans_I114T_3 | NEUEY478NZP | Male | 49 | 49 |
| Ans_A5T_4 | NEUHK991AVP | Male | 46 | 47 |
| Ans_I114T_5 | NEUTA057AF6 | Male | 63 | 65 |
| Ans_I113T_6 | NEUWM344ZLM | Female | 39 | 52 |
| Ans_I114T_7 | NEUXP289KRC | Female | 39 | 56 |
| Ans_A5T_8 | NEUZF321EW4 | Male | 46 | 47 |
| Ans_CTRL_1 | NEUAJ025JC3 | Female | 49 | |
| Ans_CTRL_2 | NEUAJ928PAA | Female | 58 | |
| Ans_CTRL_3 | NEUCA748GF2 | Female | 28 | |
| Ans_CTRL_4 | NEUCV136DHM | Male | 62 | |
| Ans_CTRL_5 | NEUCV809LL4 | Female | 50 | |
| Ans_CTRL_6 | NEUDE949BP3 | Female | 48 | |
| Ans_CTRL_7 | NEUDM126GNG | Female | 55 | |
| Ans_CTRL_8 | NEUEU392AE8 | Female | 50 | |
| Ans_CTRL_9 | NEUEY565NWT | Female | 53 | |
| Ans_CTRL_10 | NEUFE306EFY | Female | 62 | |
| Ans_CTRL_11 | NEUFZ500KDB | Female | 49 | |
| Ans_CTRL_12 | NEUFZ508VBV | Female | 43 | |
| Ans_CTRL_13 | NEUHE723FGT | Male | 64 | |
| Ans_CTRL_14 | NEUJH290RH7 | Female | 70 | |
| Ans_CTRL_15 | NEUJX341NDP | Female | 35 | |
| Ans_CTRL_16 | NEUKW131XJ2 | Female | 47 | |
| Ans_CTRL_17 | NEULL933JXY | Female | 20 | |
| Ans_CTRL_18 | NEUMN061ATZ | Male | 60 | |
| Ans_CTRL_19 | NEUNW343RXP | Male | 71 | |
| Ans_CTRL_20 | NEUPH301NNX | Male | 71 | |
| Ans_CTRL_21 | NEURJ861MMD | Female | 59 | |
| Ans_CTRL_22 | NEUUV825HYF | Female | 72 | |
| Ans_CTRL_23 | NEUVZ050YX7 | Male | 49 | |
| Ans_CTRL_24 | NEUWT164JRQ | Female | 68 | |
| Ans_CTRL_25 | NEUYM205MRL | Male | 60 | |
| Ans_CTRL_26 | NEUAA485DZL | Male | 64 | |
| Ans_CTRL_27 | NEUDA782GW3 | Female | 34 | |
| Ans_CTRL_28 | NEUDT762KUL | Female | 69 | |
| Ans_CTRL_29 | NEUEB210XRC | Male | 75 | |
| Ans_CTRL_30 | NEUFL733GX5 | Female | 58 | |
| Ans_CTRL_31 | NEUHZ716BZ2 | Female | 57 | |
| Ans_CTRL_32 | NEUMA002VLD | Male | 64 | |
| Ans_CTRL_33 | NEUMF089KLV | Male | 58 | |
| Ans_CTRL_34 | NEUML507PFJ | Female | 42 | |
| Ans_CTRL_35 | NEUMT184NWC | Female | 60 | |
| Ans_CTRL_36 | NEUNC876ZB2 | Male | 36 | |
| Ans_CTRL_37 | NEUNN472ACB | Male | 27 | |
| Ans_CTRL_38 | NEUPL878MTL | Female | 61 | |
| Ans_CTRL_39 | NEUPW536ZKZ | Male | 59 | |
| Ans_CTRL_40 | NEURV546WMW | Female | 60 | |
| Ans_CTRL_41 | NEUWN092BVG | Female | 44 | |
| Ans_CTRL_42 | NEUXC258VTR | Female | 28 | |
| Ans_CTRL_43 | NEUXP955XW7 | Female | 61 | |
| Ans_CTRL_44 | NEUXW311EFC | Female | 69 | |
| Ans_CTRL_45 | NEUZL045YD3 | Female | 66 | |

1.	Kiskinis E, Sandoe J, Williams Luis A, et al. Pathways Disrupted in Human ALS Motor Neurons Identified through Genetic Correction of Mutant SOD1. *Cell Stem Cell*. 2014;14(6):781-795. doi:10.1016/j.stem.2014.03.004

2.	Wang LX, Yi F, Fu LN, et al. CRISPR/Cas9-mediated targeted gene correction in amyotrophic lateral sclerosis patient iPSCs. *Protein & Cell*. May 2017;8(5):365-378. doi:10.1007/s13238-017-0397-3

3.	Dash BP, Freischmidt A, Weishaupt JH, Hermann A. Downstream Effects of Mutations in SOD1 and TARDBP Converge on Gene Expression Impairment in Patient-Derived Motor Neurons. *Int J Mol Sci*. Aug 25 2022;23(17)doi:10.3390/ijms23179652
