## Supplementary figures and images for "Comparative Temporal Transcriptomic Analysis of *SOD1* Mutations in iPSC-Motor Neurons"

### Supplemental Fig 1

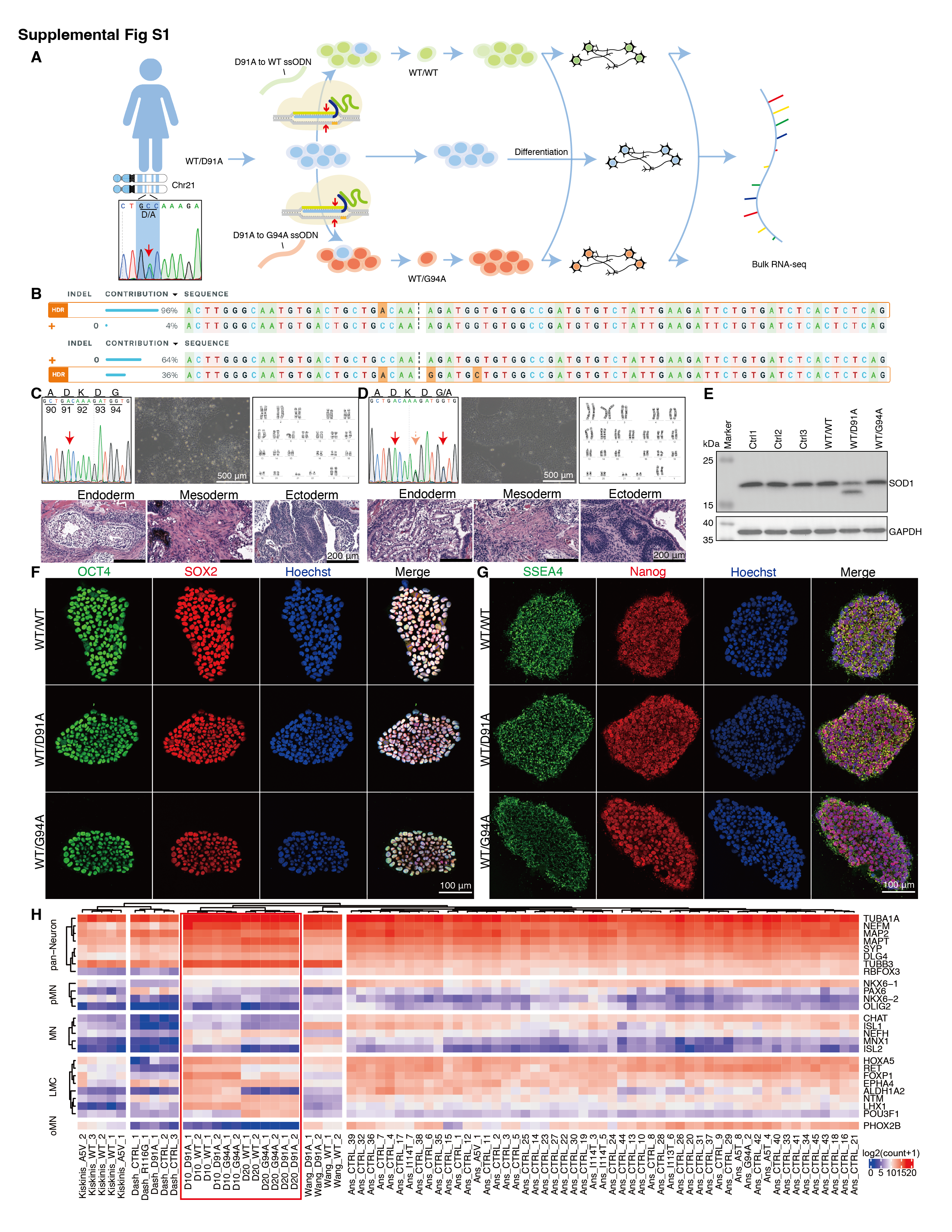

### Supplemental Fig 2

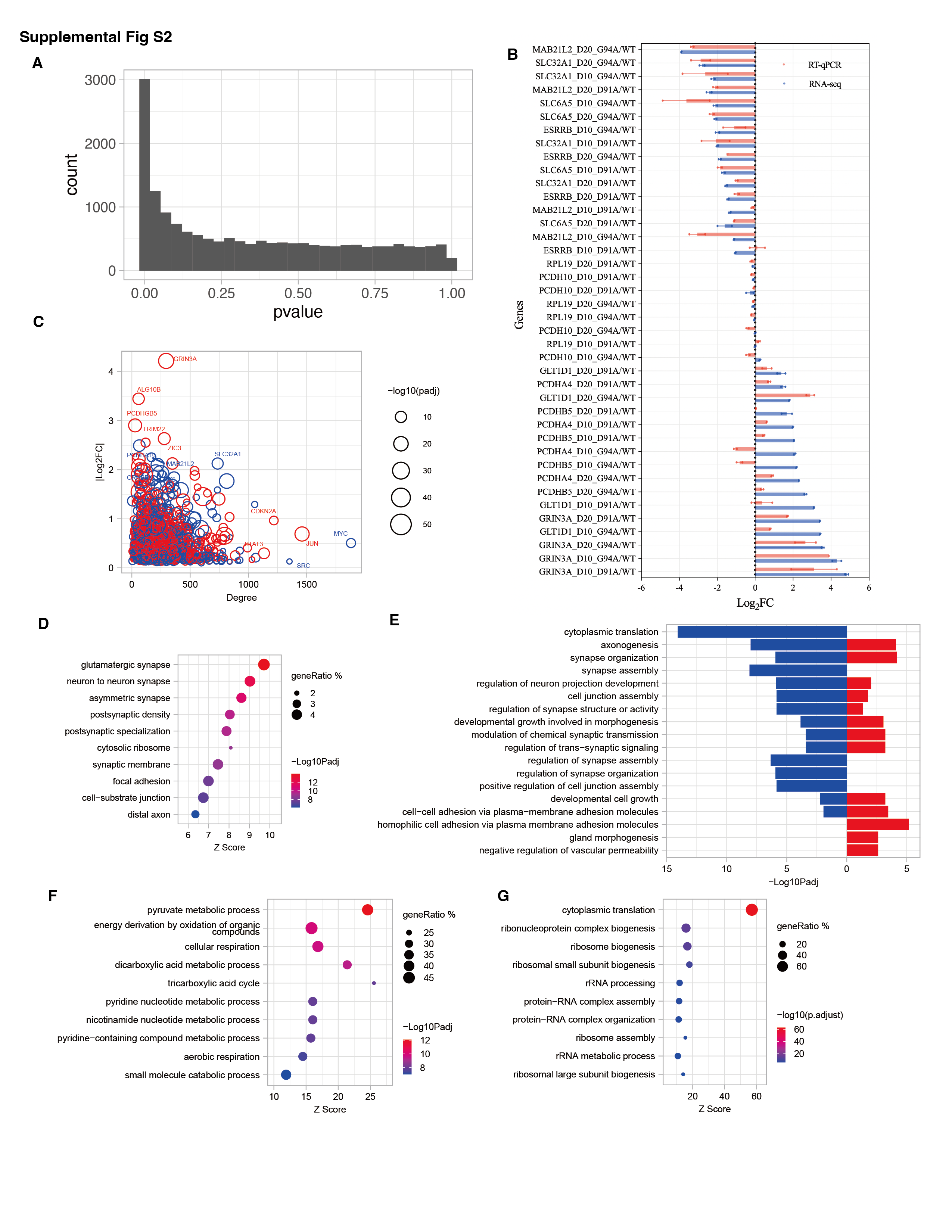
